# Adjuvant myeloperoxidase inhibition promotes B and T cell mediated control of murine melanoma metastasis

**DOI:** 10.64898/2026.11.20.768197

**Authors:** Kaitlyn M. Landreth, Adam W. Sharp, Angisha Basnet, Brian A. Boone, Mary Garland-Kledzik, Tracy W. Liu

## Abstract

Metastasis is the spread of cancer cells from primary tumors to distant organs and accounts for the majority of melanoma deaths, with a 5-year survival rate of only 30%. Despite strong evidence that myeloid cells promote metastasis, there are limited clinically effective myeloid-directed therapies. Here, we demonstrate that targeting myeloperoxidase (MPO), a myeloid-lineage restricted enzyme, significantly reduced multi-organ metastatic burden using an immunocompetent mouse model of spontaneous melanoma. Adjuvant MPO inhibition remodeled the metastatic microenvironment by a reduction in Ly6G^+^Ly6C^+^ myeloid cells and enrichment of B and T cell populations. Importantly, adjuvant MPO inhibitor-mediated metastatic control was dependent upon B, CD8^+^ T cells and CD4^+^ T cells, and rapidly reduced circulating and disseminated tumor cells. MPO inhibition markedly decreased NET release, while NET deficiency and inhibition recapitulated the anti-metastatic effects of targeting MPO, implicating NETs as a potential downstream mediator of MPO-driven metastasis. We further validated these findings in a distinct spontaneous breast cancer model where adjuvant MPO inhibition similarly reduced multi-organ metastatic burden with comparable changes within the metastatic immune microenvironment. Collectively, our studies introduce MPO as a druggable regulator of metastasis where MPO inhibition limits metastatic disease by targeting immunosuppressive myeloid cells.

## Introduction

Metastasis is the spread of cancer cells from primary tumors to distant organs and accounts for the majority of melanoma deaths, with a 5-year survival rate of only 30%.^1^ Metastasis is critically shaped by interactions between disseminating tumor cells, the immune system, and metastatic microenvironments.^2^ Increasing evidence implicates myeloid cells as key drivers of metastatic progression through suppression of anti-tumor immunity and conditioning of permissive metastatic niches.^3–6^ Clinically, accumulation of these myeloid cell populations correlate with metastatic burden and poor patient survival.^7–10^ Despite strong evidence that myeloid cells promote metastasis, there are limited clinically effective myeloid-directed therapies. Thus, defining druggable pathways that sustain myeloid cell immunosuppression and promotion of metastasis remains a critical unmet need.

Among the mechanisms employed by myeloid cells, excessive production of reactive oxygen species (ROS) is a mediator of immune suppression and metastatic progression.^11–14^ The dominant source of these oxidants in myeloid cells is myeloperoxidase (MPO), a myeloid-lineage restricted enzyme.^15^ MPO is the most abundant protein in neutrophil azurophilic granules, is approximately 1% of monocyte protein, and also found in myeloid-derived suppressor cells (MDSCs) and macrophages.^16–20^ MPO primarily catalyzes the production of hypochlorous acid that mediate antimicrobial activities^15,20–22^ while MPO-dependent ROS production have been shown to suppress adaptive immune responses.^19,23–26^ Additionally, MPO has been shown to promote neutrophil extracellular trap (NET) formation^27–29^ which been implicated is promoting metastasis.^30–33^ However, the contribution of MPO, itself, to metastatic progression remains unknown. Here, we report the effect and mechanism of adjuvant MPO inhibition which reduces multi-organ metastatic melanoma burden in a B and T cell dependent manner with additional pharmacological evidence of the efficacy of adjuvant MPO inhibition in a breast cancer metastasis model.

## Results

### MPO deficiency and inhibition reduces multi-organ melanoma metastatic burden

We previously demonstrated that MPO was elevated in melanoma compared to normal skin.^34^ Analysis of the TCGA-SCKM dataset further confirmed a significantly higher MPO expression in metastatic melanoma compared to primary tumors (Fig. S1A). The impact of MPO on metastatic burden was evaluated in a syngeneic, immunocompetent spontaneous melanoma model using a B16F10 cell line stably expressing click beetle green luciferase and enhanced green fluorescent protein (B16F10 CBG-GFP).^35,36^ Primary subcutaneous tumors were resected 18 days after subcutaneous injection of B16F10 CBG-GFP cells, with mice developing distant metastases in the lungs, bone and inguinal lymph nodes by week 6, mimicking the clinical patterns of metastatic spread in melanoma. Metastasis was compared in wild type C57BL/6J (WT) and age-matched syngeneic MPO-deficient (*MPO^-/-^*, C57BL/6J background^37^) mice. To evaluate the effects of pharmacologic MPO inhibition, WT mice were treated with the MPO-specific inhibitor mitiperstat.^38,39^ Adjuvant MPO inhibitor treatment began on day 19, one day post subcutaneous tumor resection, and consisted of daily intraperitoneal (IP) injections of mitiperstat for 21 days (Fig. 1A). Both genetic MPO deficiency and pharmacologic MPO inhibition significantly reduced multi-organ B16F10 metastatic burden, as quantified by bioluminescence imaging (Fig. 1B – D). All WT mice developed metastases at one or more distant sites. In contrast, 39.1% of MPO-deficient mice (9/23) and 57.1% of mitiperstat-treated WT mice (12/21) exhibited no bioluminescence detectable metastases at any of the metastatic sites (Fig. 1E). Bioluminescence detectable metastases were observed in 100%, 95.7%, and 95.7% of WT mice in the lung, bone, and lymph nodes, respectively (Fig. S1B). In contrast, only 43.5%, 43.5%, and 30.4% of MPO-deficient mice and only 4.8%, 23.8%, and 28.6% of mitiperstat treated WT mice developed bioluminescence detectable metastases in the lung, bone, and lymph nodes, respectively (Fig. S1B). This significant decrease in metastasis from MPO deficiency and inhibition was further confirmed by cresyl violet staining of lung tissue (Fig. 1F). No sex-dependent differences in metastatic burden were observed in WT, *MPO^-/-^* or MPO inhibitor treated WT mice (Fig. S1C). Mice tolerated MPO inhibitor treatment well, with no significant changes in body weight (Fig. S1D). Comparing primary subcutaneous tumor growth, MPO deficiency resulted in significantly decreased tumor volumes at days 11 and 14 after B16F10 cell injection compared to WT mice; however, this difference was no longer evident by day 18 (Fig. S1E). Because MPO inhibitor treatment was initiated after subcutaneous tumor resection, the effects of mitiperstat on subcutaneous tumor growth were not assessed. Taken together, these data suggest that targeting MPO markedly reduced multi-organ metastatic disease.

**Figure 1.**
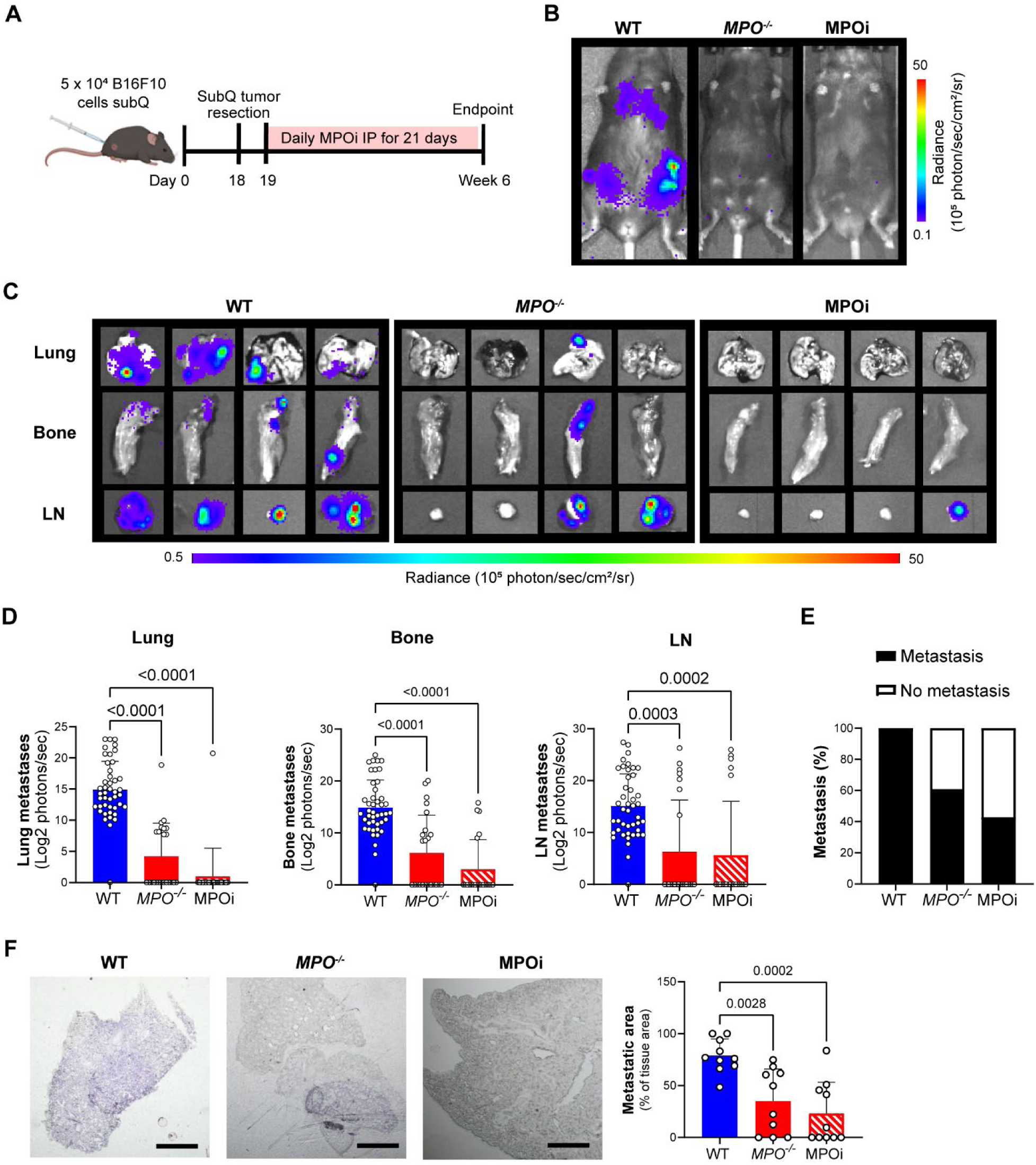
MPO deficiency and inhibition significantly reduces multi-organ metastatic melanoma burden. (A) Schematic of the spontaneous B16F10 metastasis model and MPO inhibitor, mitiperstat, (MPOi) treatment consisting of daily intraperitoneal injections for 21 days beginning on day 19. Representative (B) *in vivo* and (C) *ex vivo* B16F10 CBG-GFP bioluminescence imaging. Corresponding (D) *ex vivo* bioluminescence quantification of metastatic burden in the lung, bone and inguinal lymph nodes (LN) and (E) percentage of mice with detectable metastasis at experimental endpoint in WT, *MPO^-/-^* and MPOi mitiperstat treated WT mice (n = 44 WT, n = 23 *MPO^-/-^*, n = 21 MPOi; equal sex). (F) Representative images of cresyl violet staining and corresponding quantification of metastatic area in of lung tissues (n = 10 mice/group; equal sex). Data shown as mean ± SD; statistical significance of differences in tumor volume calculated using a two-way ANOVA followed by Tukey’s multiple comparisons test (p values shown on graphs).

### MPO deficiency and inhibition remodels the metastatic immune microenvironment

To determine how MPO influences the metastatic immune microenvironment, we performed flow cytometry analysis of metastatic tissues at the experimental endpoint. Both MPO deficiency and inhibition significantly reduced the frequency of Ly6G^+^Ly6C^+^ myeloid cells across all metastatic sites with over a 10-fold decrease compared to WT controls (Fig. 2A). A significant decrease in CD11b^+^Ly6G^+^ neutrophils were observed in *MPO^-/-^*and mitiperstat-treated WT mice in the lungs and lymph nodes while CD11b^+^Ly6C^+^ monocytes showed metastatic site-specific changes (Fig. S2A&B). These changes were accompanied by significantly increased frequencies of CD8⁺ T cells in the bone and inguinal lymph nodes (approximately a 2 – 5-fold increase) and B cells in the lungs and lymph nodes (approximately a 3 – 6-fold increase) in *MPO^-/-^* and MPO inhibitor treated WT mice compared to WT controls (Fig. 2B&C). Phenotypic characterization of CD8⁺ T cells demonstrated increased frequencies of TRP2⁺ tumor-specific, CD107a⁺ cytotoxic, CD44⁺ antigen-experienced, and CD44⁺CD62L⁺ central memory CD8⁺ T cells across all metastatic sites in both *MPO^-/-^*and MPO inhibitor treated WT mice compared to WT controls (Fig. 2D – G). Notably, TRP2⁺ and CD107a⁺ CD8⁺ T cells were increased by more than 18-fold in the inguinal lymph nodes, and CD44⁺ CD8⁺ T cells were increased by more than 20-fold in the lungs from MPO deficiency and inhibition. Additional effector and activation associated phenotypes exhibited site-specific changes from targeting MPO, including increased Granzyme B^+^ and CD25^+^ CD8^+^ T cell frequencies in selected metastatic tissues (Fig. S2C&D). Other immune cell populations including dendritic cells and macrophages showed metastatic site-specific changes, whereas natural killer (NK) cell frequencies were largely unchanged. (Fig. S2E – G). CD4^+^ T cells also exhibited changes in abundance and phenotype. MPO deficiency and inhibition significantly increased CD4^+^ T cell frequencies in the lymph nodes whereas MPO inhibition decreased these cells in the lungs (Fig. S2H). Multiple CD4^+^ T cell populations expressing TRP2⁺, CD107a⁺, CD44⁺, CD44⁺CD62L⁺, Granzyme B^+^ or CD25^+^ were significantly enriched in the lungs and/or lymph nodes from MPO deficiency and inhibition compared to WT controls (Fig. S2I – N). Collectively, these data suggest that MPO deficiency or inhibition remodels the metastatic immune tumor microenvironment resulting in a reduction of myeloid populations and enrichment of B cell frequencies and T cell tumor-specific, activated and effector associated phenotypes within metastatic tissues.

**Figure 2.**
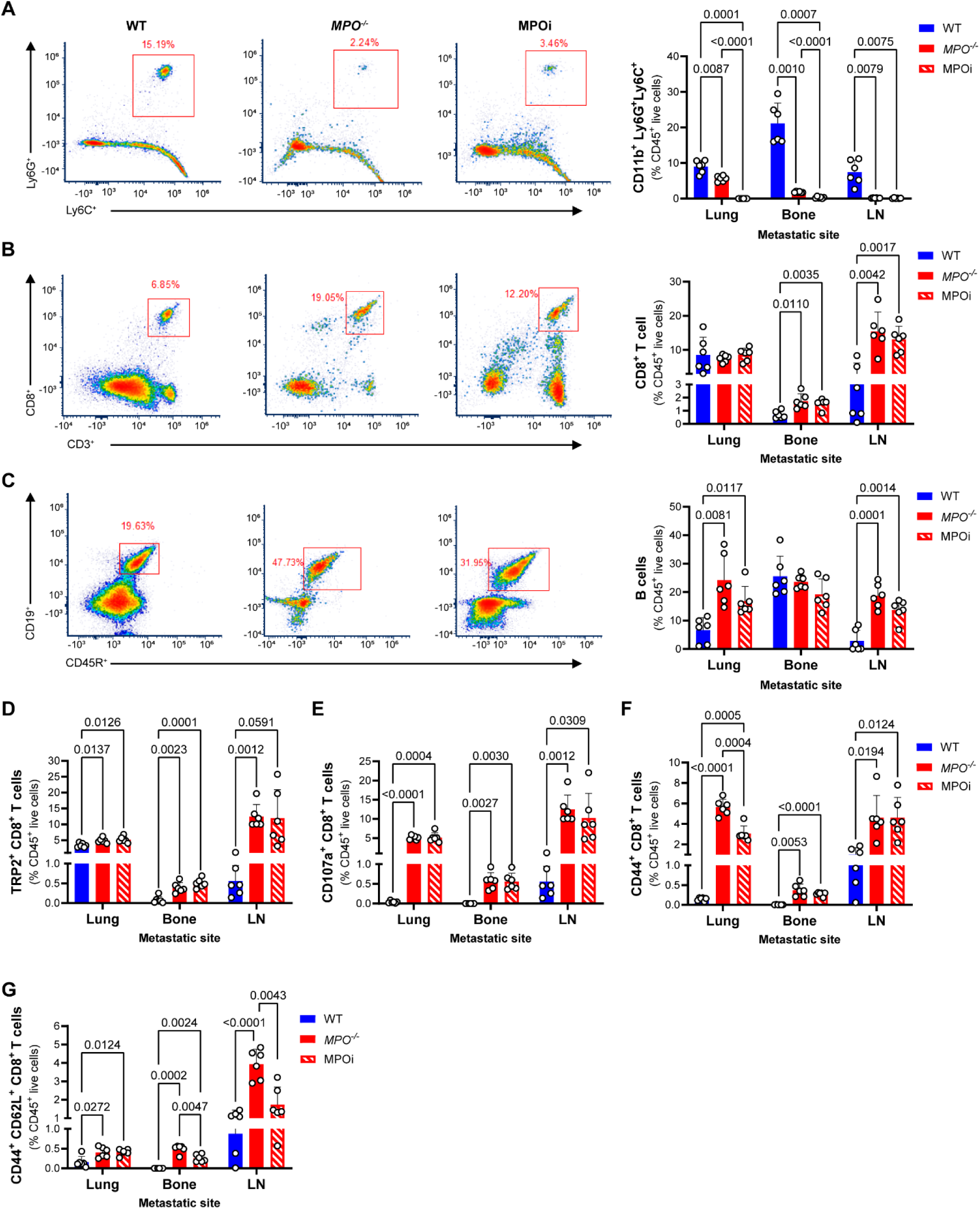
MPO deficiency and inhibition significantly remodels the metastatic immune microenvironment. Representative flow cytometry plots and corresponding quantification of frequencies (%) of (A) Ly6G^+^Ly6C^+^ myeloid cells, (B) B cells and (C) CD8^+^ T cells in the lungs, bone, and lymph nodes (LN) from spontaneous B16F10 metastatic WT, *MPO^-/-^* and MPOi mitiperstat treated WT mice at experimental endpoint. MPO deficiency and inhibition with mitiperstat significant increases the frequencies (%) of (D) TRP2^+^, (E) CD107a^+^, (F) CD44^+^ and (G) CD44^+^CD62L^+^ CD8^+^ T cells across all metastatic sites compared to WT mice. Data shown as mean ± SD (n = 6 mice/group, equal sex); statistical significance of differences in immune cell frequencies calculated using a two-way ANOVA followed by Tukey’s multiple comparisons test (p values shown on graphs).

### Targeting MPO reduces metastatic burden through inhibition of NETs

Given that MPO is a major producer of ROS, we evaluated whether its pro-metastatic effects were ROS-dependent. Upstream of MPO in the oxidative burst pathway is NADPH oxidase 2 (NOX2), which generates the hydrogen peroxide used by MPO to primarily produce hypochlorous acid. Metastatic burden was compared in WT and age-matched syngeneic NOX2-deficient (*NOX2^-/-^*) mice. NOX2-deficiency did not reproduce the multi-organ reduction in metastatic burden observed with targeting MPO where100% of *NOX2^-/-^* mice had detectable bioluminescence metastases at one distant site (Fig. S3A). While NOX2-deficiency resulted in significant reduction of metastatic burden within the bone (Fig. S3B), no differences were observed in the lungs or inguinal lymph nodes (Fig. S3C&D). This suggests that the reduction in ROS is not sufficient to explain the anti-metastatic effects of targeting MPO. Given that MPO has also been shown to promote the formation of NETs, the impact of NETs on metastatic burden was also evaluated. Metastatic burden was compared in WT and age-matched syngeneic peptidyl arginine deiminase 4-deficient mice (*PAD4^-/-^*, C57BL/6J background)^40^ which have impaired PAD4-dependent NET formation. The effects of hydroxychloroquine (HCQ), shown to be a NETs inhibitor^41–43^, was also evaluated in WT mice. Similar to MPO inhibition, HCQ treatment began one day post primary tumor resection on day 19 and consisted of 0.5 mg/mL HCQ dissolved in drinking water for 21 days. Similar to targeting MPO, both NETs deficiency and inhibition resulted in significant reduction of B16F10 metastatic burden measured at all metastatic sites by *ex vivo* bioluminescence (Fig. 3A). 40% of PAD4-deficient mice (6/15) and 58.6% of HCQ-treated WT mice (17/29) exhibited no detectable bioluminescence metastases at any of the metastatic sites (Fig. 3B). Bioluminescence detectable metastases were observed in 40.0%, 13.3%, and 46.7% of *PAD4^-/-^*mice and 34.5%, 17.2%, and 13.8% of HCQ-treated WT mice in the lungs, bone, and inguinal lymph nodes, respectively (Fig. S4A). Male *PAD4^-/-^* mice developed significantly higher lung metastatic burden compared to female mice but no other sex-dependent differences in metastatic burden were observed in *PAD4^-/-^*or HCQ treated WT mice (Fig. S4B&C). Mice tolerated HCQ treatment well, with no significant changes in body weight (Fig. S4D). Comparing primary subcutaneous tumor growth, PAD4 deficiency led to smaller tumor volumes at days 11 and 14 after B16F10 cell injection; however, by day 18, when primary tumors were resected, no difference in tumor size was observed (Fig. S4E). Because HCQ treatment was initiated after primary tumor resection, the effects of HCQ on subcutaneous tumor growth were not assessed.

**Figure 3.**
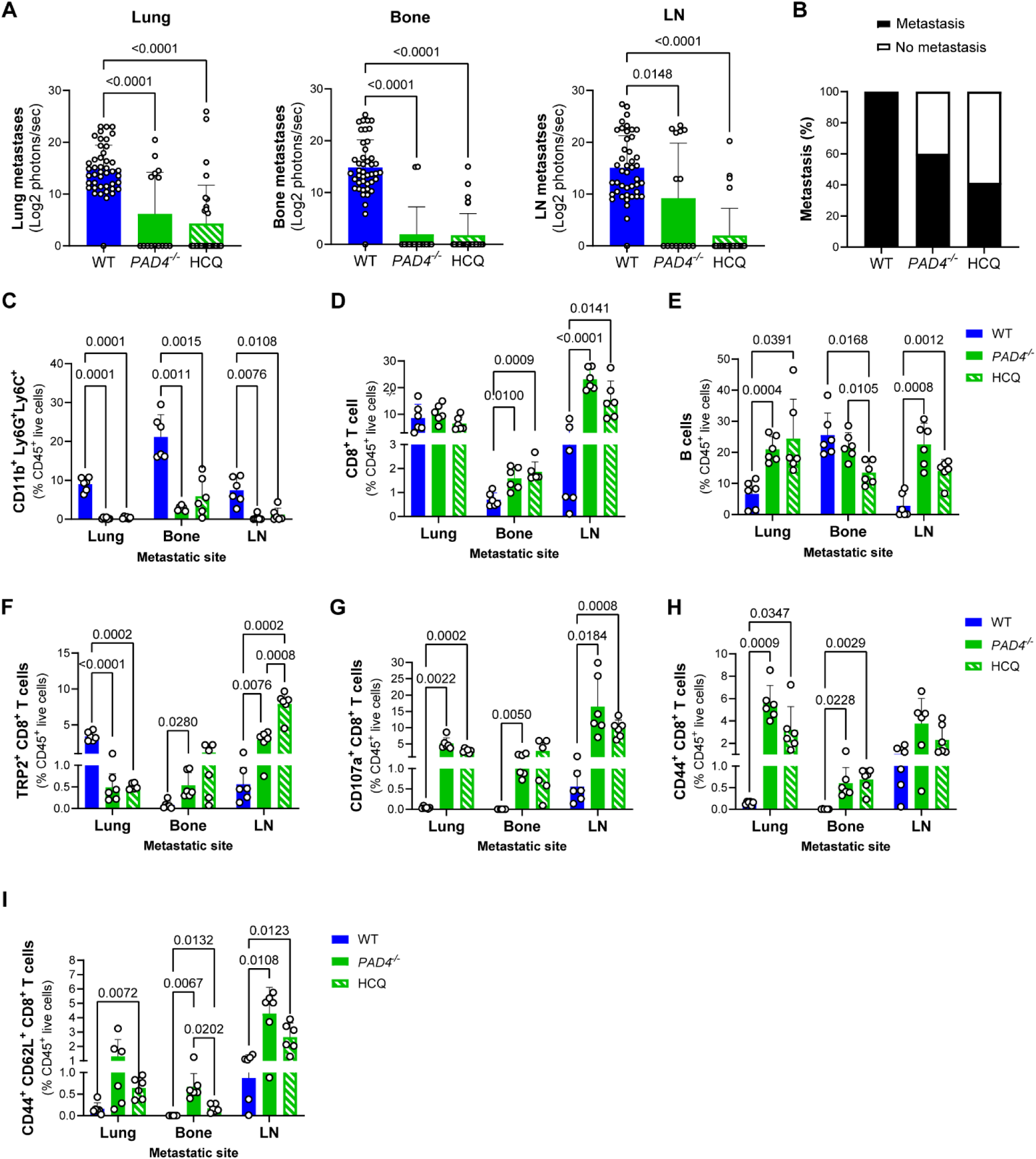
NET deficiency and inhibition phenocopies targeting MPO. (A) *Ex vivo* bioluminescence quantification of metastatic burden in the lung, bone and lymph nodes (LN) and (B) percentage of mice with detectable metastasis at experimental endpoint in WT, *PAD4^-/-^* and hydroxychloroquine treated WT (HCQ) mice (n = 44 WT same dataset as shown in figure 2, n = 15 *PAD4^-/-^*, n = 29 HCQ; equal sex). Flow cytometry quantification of frequencies (%) of (C) Ly6G^+^Ly6C^+^ myeloid cells, (D) B cells, (E) CD8^+^ T as well as CD8^+^ T cell subsets expressing (F) TRP2^+^, (G) CD107a^+^, (H) CD44^+^ and (I) CD44^+^CD62L^+^ metastatic tissues from spontaneous B16F10 metastatic WT, *PAD4^-/-^*and HCQ treated WT mice at experimental endpoint (n = 6 mice/group, equal sex; WT dataset same as shown in figure 3). Data shown as mean ± SD; statistical significance of differences calculated using a two-way ANOVA followed by Tukey’s multiple comparisons test (p values shown on graphs).

To investigate whether PAD4 deficiency and HCQ treatment reproduced the immune alterations observed from MPO targeting, we profiled metastatic tissues of *PAD4^-/-^* and HCQ-treated WT mice. Similar to MPO deficiency and inhibition, both PAD4 deficiency and HCQ treatment significantly reduced the Ly6G^+^Ly6C^+^ myeloid cell frequencies across all metastatic sites (Fig. 3C). A significant increase in the frequencies of CD8^+^ T cells in the bone and lymph nodes and B cells in the lung and lymph nodes of *PAD4^-/-^* and HCQ treated WT mice compared to WT controls paralleled the changes observed from targeting MPO (Fig. 3D). However, HCQ treatment additionally reduced B cell frequencies in the bone relative to WT controls and *PAD4^-/-^* mice (Fig. 3E). Phenotypic analysis of CD8^+^ T cell frequencies revealed overlapping but non-identical responses compared to targeting MPO. Both PAD4-deficiency and HCQ treatment increased TRP2^+^ CD8^+^ T cells in the lymph nodes, were only increased in the bone of *PAD4^-/-^*compared to WT mice and were decreased in the lungs of *PAD4^-/-^* and HCQ treated WT mice compared to WT controls (Fig. 3F). A significant increase in the frequencies of CD107a^+^ CD8^+^ T cells in the lungs and lymph nodes, CD44^+^ CD8^+^ T cells in the lung and bones, CD44^+^CD62L^+^ CD8^+^ T cells in the bone and lymph nodes of *PAD4^-/-^* and HCQ treated WT mice compared to WT controls (Fig. 3G – I). Granzyme B^+^ CD8^+^ T cell frequency was not significantly altered whereas a significant increase in CD25^+^CD8^+^ T cell frequency occurred in the lungs and lymph nodes of *PAD4^-/-^* and HCQ treated WT mice compared to WT controls (Fig. S5A&B). Other immune cell populations exhibited site-specific changes from PAD4-deficiency and HCQ treatment (Fig. S5C - G). Phenotypic analysis of CD4^+^ T cells revealed additional site-specific enrichment of tumor-specific, cytotoxic, antigen-experienced, central memory, and effector-associated populations of CD4^+^ T cells, particularly in the lungs and lymph nodes in *PAD4^-/-^* and HCQ treated WT mice compared to WT controls (Fig. S5H – N). These observations demonstrate that PAD4-deficiency and HCQ treatment reproduce several of the immune alterations associated with MPO targeting, including reduced Ly6G^+^Ly6C^+^ myeloid cell frequencies and enrichment of B cells and selected CD8^+^ T cell phenotypes. Taken together, these data support NETs as a downstream mechanism by which MPO promotes metastasis.

### Targeting MPO reduces the immunosuppressive activity of Gr-1+ myeloid cells

Given the significant decrease in Ly6G^+^Ly6C^+^ myeloid cells from MPO deficiency and inhibition at metastatic sites, we isolated Gr-1+ myeloid cells from the lungs of metastatic mice at experimental endpoint to evaluate how MPO impacted their function. Aligning with our flow cytometry data, we observe a significant decrease in lung-isolated Gr-1+ myeloid cells when MPO or NETs were deficient or inhibited (Fig. S6A). A significant decrease in MPO activity was observed in Gr-1+ myeloid cells from MPO or NET deficient or inhibited mice relative to WT cells, measured by luminol bioluminescence (Fig. 4A). ROS levels were reduced in Gr-1+ myeloid cells isolated from *MPO^-/-^*, *PAD4^-/-^*and HCQ treated mice but not mitiperstat treated mice compared to WT controls, measured using L-012 (Fig. 4B). Significantly reduced NET release in Gr-1+ myeloid cells was observed when MPO or NETs were deficient or inhibited compared to WT measured using a Pico green assay (Fig. 4C). To determine whether these changes altered the immunosuppressive function, equal numbers of viable Gr-1+ myeloid cells were co-cultured with CD8^+^ T cells isolated from the spleen of healthy (tumor-free) WT mice. WT myeloid cells markedly suppressed CD8^+^ T cell proliferation, whereas this suppressive activity was significantly attenuated when MPO or NETs were deficient or inhibited (Fig. 4D). This loss of suppressive activity was accompanied by increased TNFα levels in Gr-1+ myeloid cell and CD8^+^ T cell co-culture supernatants (Fig. 4E). Similarly, equal number of Gr-1+ myeloid cells were co-cultured with B cells isolated from the spleen of healthy WT mice where a significant increase in B cell proliferation was observed with MPO or NETs deficiency and inhibition relative to WT cells (Fig. 4F). This loss of suppressive activity on B cells was accompanied by significantly decreased interleukin-10 (IL-10) and IgM levels (Fig. 4G&H) while no difference was observed in IgA or IgG levels (Fig. S6B&C) in Gr-1+ myeloid cell and B cell co-culture supernatants. Cytokine profiling also demonstrated that MPO deficiency and inhibition resulted in a significant decrease in secreted interleukin-1 alpha (IL-1α) levels in Gr-1+ myeloid cells when MPO or NETs were deficient or inhibited (Fig. S6D).

**Figure 4.**
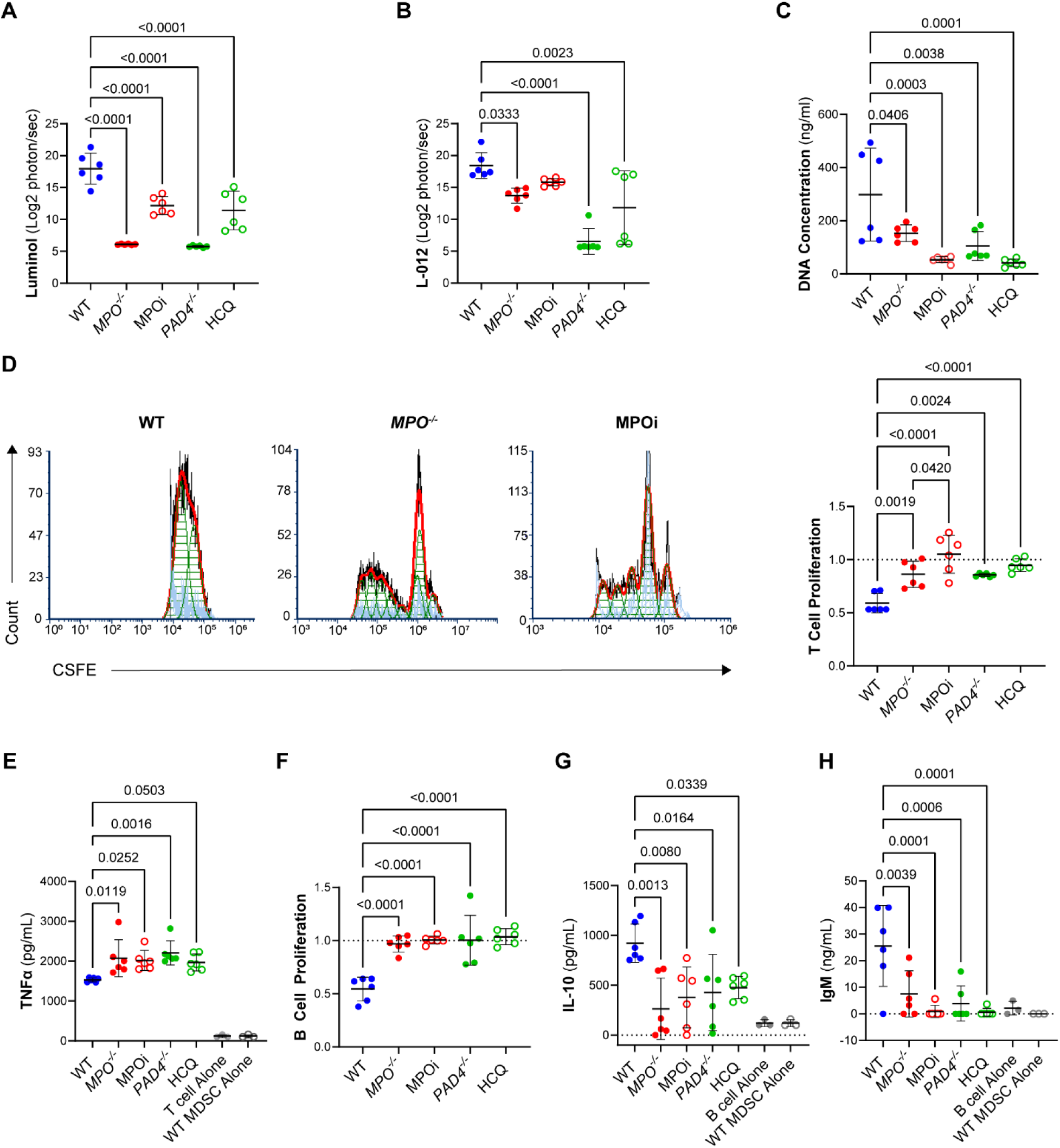
MPO deficiency and inhibition reduces the immunosuppressive activity of Gr-1+ myeloid cells. Lung-isolated Gr-1+ myeloid cells from spontaneous B16F10 metastatic *MPO^-/-^*or mitiperstat treated (MPOi) WT mice at experimental endpoint show reduced (A) MPO activity measured by luminol, (B) ROS levels measured by L-012, and (C) NETs measured using a Pico Green assay. (D) Representative T cell proliferation plots and corresponding quantification of spleen-isolated CD8^+^ T cells from healthy WT mice co-cultured with Gr-1+ myeloid cells at 72 h assessed by CFSE dilution (normalized to CD3, CD28 activated CD8^+^ T cells alone) and (E) TNFα levels in the supernatant. (F) B cell proliferation of spleen-isolated B cells from healthy WT mice co-cultured with Gr-1+ myeloid cells at 72 h assessed by CFSE dilution (normalized to IgM activated B cells) and (G) IL-10 and (H) IgM levels in the supernatant. Data shown as mean ± SD (n = 6 mice/group, equal sex); statistical significance of differences calculated using a two-way ANOVA followed by Tukey’s multiple comparisons test (p values shown on graphs).

To determine whether mitiperstat or HCQ treatment impaired the ability of Gr-1+ myeloid cells to promote metastatic growth, we isolated splenic Gr-1+ myeloid cells from metastatic WT controls, mitiperstat or HCQ treated WT mice at week 6 and subcutaneously co-injected them with B16F10 CBG-GFP cells at a 1:1 ratio into WT recipient mice (Fig. S7A). The addition of WT Gr-1+ myeloid cells significantly enhanced subcutaneous tumor growth compared to co-injection of myeloid cells mitiperstat or HCQ treated mice or injection of tumor cells alone (Fig. S7B - D). However, following resection of subcutaneous tumors, no differences in metastatic burden was observed at experimental endpoint (Fig. S7E). These findings suggest that MPO inhibition and HCQ treatment diminish the primary tumor-promoting activity of Gr-1^+^ myeloid cells without subsequent detectable differences in metastatic burden using this co-injection model. We also selectively depleted Ly6G⁺ myeloid cells to evaluate whether the effects of MPO were specific to Ly6G^+^ myeloid cells. Ly6G⁺ myeloid cell depletion resulted in only a partial reduction of bone and lymph node metastases and did not reduce lung metastasis (Fig. S8), failing to phenocopy MPO targeting suggesting that MPO inhibition exerts broader immunomodulatory effects. Taken together, these data suggest that MPO regulates the immunosuppressive activity of Gr-1+ myeloid cells and hinders T and B cell proliferation and activation.

### Anti-metastatic efficacy of MPO inhibition requires B and T cells

Given that MPO inhibition enriched B and T cell frequencies within metastatic tissues and attenuated Gr-1+ myeloid cell-mediated immune suppression, we next determined whether these B and T cells were required for the anti-metastatic effects of MPO inhibition. The contribution of B cells was evaluated by systemic depletion using an anti-CD20 antibody in WT mice treated with or without mitiperstat. IgG treated WT mice, with or without MPO inhibition were included controls. While B cell depletion alone did not significantly alter metastatic burden compared to IgG treated WT controls, B cell depletion in mitiperstat treated mice significantly increased lung and bone metastatic burden, effectively reversing the anti-metastatic effects of MPO inhibition and restoring metastatic burden to levels comparable to those observed in IgG treated controls (Fig. 5A). We noted that in the IgG treated controls, MPO inhibition significantly reduced lung and bone metastasis, but the reduction in lymph node metastasis did not reach statistical significance. We next investigated whether CD8^+^ and CD4^+^ T cells were similarly involved in the anti-metastatic effects of MPO inhibition. Individual depletion of CD8^+^ or CD4^+^ T cells in mitiperstat treated WT mice significantly increased metastatic burden in the lung and bones compared to mitiperstat treated IgG controls, reversing the reduced metastatic burden associated with MPO inhibition (Fig. 5B&C). To determine whether these immune cell requirements observed following MPO inhibition extended to HCQ-mediated metastatic control, we performed depletion experiments in HCQ treated WT mice. Depletion of B cells, CD8^+^ T cells, or CD4^+^ T cells significantly increased metastatic burden compared to HCQ treated IgG controls (Fig. S9A – C), indicating that these adaptive immune populations also contribute to metastatic control during HCQ treatment. These findings support overlapping adaptive immune requirements following MPO inhibition and HCQ treatment. Depletion of NK cells did not significantly alter metastatic burden in IgG, mitiperstat or HCQ treated mice (Fig. S9D & E). Flow cytometry of spleens confirmed efficient depletion of B, CD8^+^ T, CD4^+^ T, and NK cells with each targeted population comprising 1% or less of immune cells following administration of the corresponding depleting antibody compared to IgG controls (Fig. S9F – G). Together, these findings demonstrate that the anti-metastatic effects of MPO inhibition were dependent upon B cells, CD8^+^ T cells and CD4^+^ T cells where their depletion disrupted mitiperstat-mediated metastatic control.

**Figure 5.**
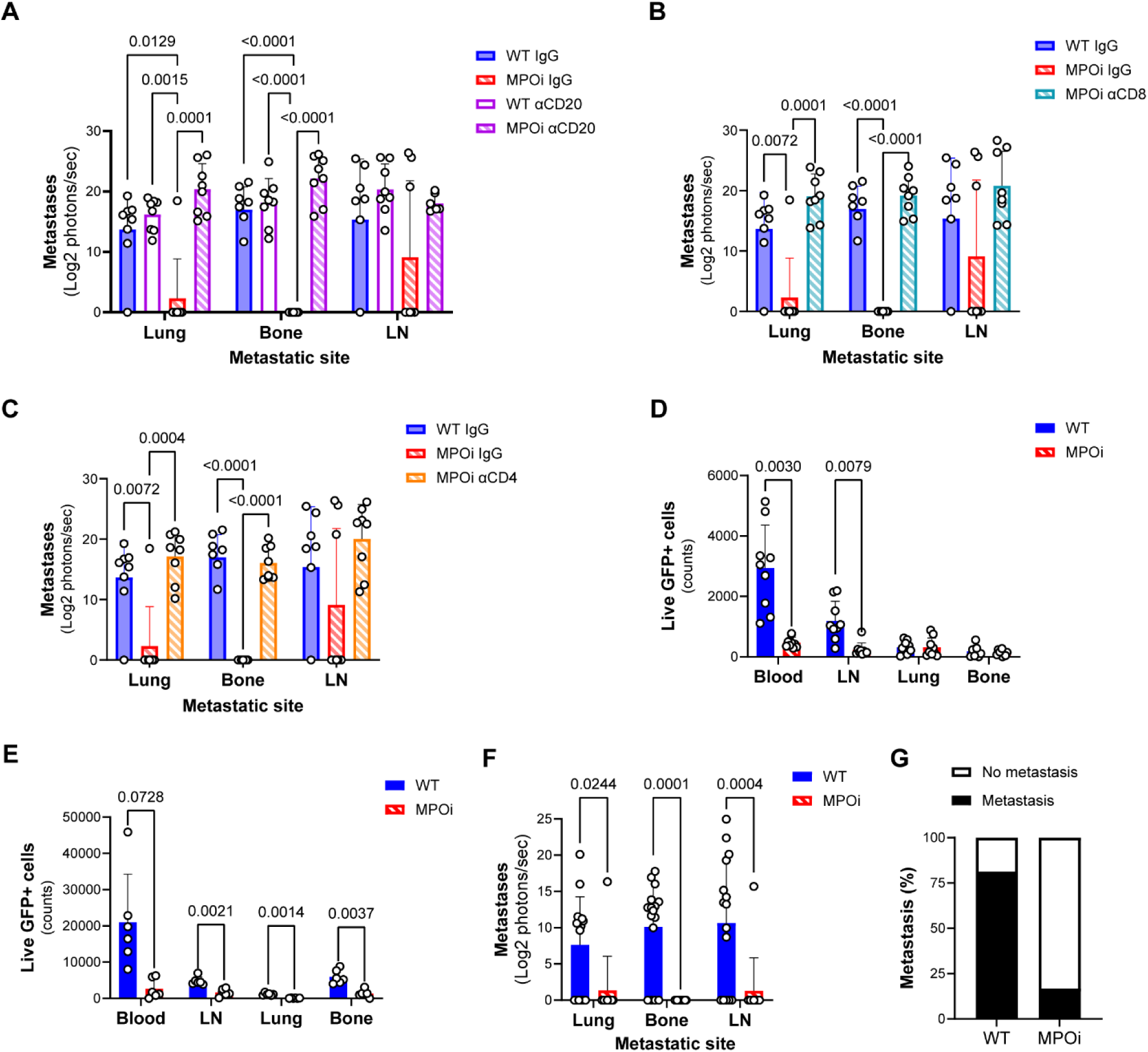
Anti-metastatic efficacy of MPO inhibition requires B and T cells while MPO inhibition reduces circulating and disseminated tumor cells. *Ex vivo* bioluminescence quantification of metastatic burden in spontaneous B16F10 metastatic WT and MPO inhibitor mitiperstat treated (MPOi) mice in the lungs, bone and inguinal lymph nodes (LN) at experimental endpoint following selective depletion of (A) B cells using an anti-CD20 (αCD20) antibody, (B) CD8+ T cells using an anti-CD8α antibody (αCD8), and (C) CD4+ T cells using an anti-CD4 antibody (αCD4). IgG treated WT and MPOi mice included controls. Data shown as mean ± SD (n=8 mice/group; equal sex). Flow cytometry quantification of GFP+ B16F10 cells following (D) a single dose of mitiperstat at day 19 and (E) 10 days of mitiperstat treatment at day 28 in the blood, LN, lungs and bone of spontaneous B16F10 metastatic WT and MPOi mice (n = 6 mice/group; equal sex). (F) *Ex vivo* bioluminescence quantification of metastatic burden and (G) percentage of mice with detectable metastasis at day 28 in WT and MPOi mice (n = 16 WT, n = 12 MPOi; equal sex). Data shown as mean ± SD (n = 6 mice/group, equal sex); statistical significance of differences calculated using a two-way ANOVA followed by Tukey’s multiple comparisons test or Šídák’s multiple comparisons test (p values shown on graphs).

### MPO inhibition reduces circulating and disseminated tumor cells and induces early immune remodeling at metastatic sites

Having demonstrated that MPO contributes to metastatic outgrowth, we next investigated whether MPO also regulates tumor cell dissemination. A single dose of mitiperstat significantly reduced circulating tumor cells in the blood and disseminated tumor cells in the inguinal lymph nodes within 24h, as quantified by GFP expression using flow cytometry (Fig. 5D). Sustained MPO inhibitor treatment for 10 days continued to reduce tumor cells in circulation, although it did not reach statistical significance, and significantly reduced tumor cells across all metastatic target sites (Fig. 5E). 10 days of mitiperstat treatment significantly decreased *ex vivo* metastatic burden compared to WT controls on day 28, preceding our established 6 week experimental endpoint (Fig. 5F). Only 16.7% of MPO inhibitor treated WT mice while 81.3% of WT mice had detectable metastasis at one or more distant sites (Fig. 5G). *Ex vivo* bioluminescence quantification revealed a 4.8-fold increase in metastatic burden at 6 weeks compared with 4 weeks in the lungs and bone (Fig. S10A). To understand whether MPO impacted the migration of tumor cells and Gr-1+ myeloid cells, transwell migration studies were evaluated. MPO inhibition reduced B16F10 cell migration toward Gr-1+ myeloid cells (Fig. S10B) while MPO inhibition did not alter myeloid cell migration towards B16F10 cells (Fig. S10C). HCQ treatment resulted in similar reduction in tumor cells in circulation and metastatic target sites after a single dose and sustained treatment for 10 days (Fig. S10D&E). However, only a significant reduction in the metastatic burden within the bones was observed on day 28 (Fig. S10F) where 57.9% of HCQ treated mice had no detectable bioluminescent metastasis (Fig. S10G). HCQ did not alter B16F10 cell or Gr-1+ myeloid cell migration (Fig. S10H&I).

To evaluate whether immune alterations within the metastatic microenvironment also emerged early, we performed flow cytometry analysis on day 28 in spontaneous metastatic WT and MPO inhibitor treated WT mice. Following 10 days of MPO inhibitor treatment, a significant reduction in Ly6G^+^Ly6C^+^ at all metastatic sites was observed while a significant increase in the frequencies of B cells in the lungs and CD8^+^ T cells in the inguinal lymph nodes was observed compared to WT controls (Fig. S11A – C). Phenotypic analysis revealed tissue-specific changes in CD8^+^ T cell subsets following MPO inhibition, characterized by increased frequencies of Granzyme B^+^ CD8^+^ T cells in the lymph nodes and CD44^+^ CD8^+^ T cells in the bone, accompanied by decreased frequencies of central memory and activated CD8^+^ T cells in the bone compared to untreated WT controls (Fig. S11D – G). No other differences in CD8⁺ T cell subsets were observed at the 4-week timepoint (Fig. S11H&I). MPO inhibitor treatment for 10 days also resulted in decreased frequencies of monocytes in the lymph nodes, NK cells in the lungs and bone, and macrophages in the bone with no changes in Ly6G^+^ neutrophils or dendritic cells compared to WT controls (Fig. S11J – N). Similarly, 10 days of MPO inhibitor treatment decreased the frequencies of CD4^+^ T cells in the lungs and bone while increasing them in the lymph nodes (Fig. S11O). Further phenotypic characterization revealed a reduction in CD4^+^ T subsets within the bone, with the exception of increased Granzyme B^+^ CD4^+^ T cell frequencies in the lymph nodes compared to WT controls (Fig. S11P – U). We also evaluated the immune composition of metastatic tissues following 10 days of NET inhibition with HCQ, which recapitulated several changes observed with MPO inhibition. HCQ treatment reduced Ly6G^+^Ly6C^+^ myeloid cell frequencies in the bone and lymph nodes with an increase in B cells in the lungs and CD8^+^ T cells in the lymph nodes (Fig. S12A – C). Generally, the frequencies of CD8^+^ and CD4^+^ T cell subsets were increased in the lymph nodes, whereas these populations were unchanged or decreased in the lungs and bone with site-specific decreases in other immune populations (Fig. S12D – U). Collectively, these findings demonstrate that adjuvant MPO inhibition reduced circulating and disseminated tumor cells and early metastatic burden, accompanied by immune remodeling characterized by reduced Ly6G^+^Ly6C^+^ myeloid cell and increased B and CD8^+^ T cell frequencies at select metastatic sites.

### MPO inhibition significantly reduces multi-organ breast cancer metastasis

To evaluate whether the effects of MPO inhibition extend beyond the B16F0 melanoma model, we used the 4T-1 triple negative breast cancer spontaneous metastasis model. These 4T-1 cells stably express firefly luciferase and enhanced GFP. After resection of the primary mammary fat pad tumor at 18 days post tumor cell implantation, mice developed metastasis in the lungs, liver, lymph nodes and bones by week 7. The same daily adjuvant MPO inhibitor, mitiperstat, treatment occurred one day after primary tumor resection which lasted for 21 days (Fig. 6A). Similar to the melanoma metastasis model, MPO inhibition significantly decreased metastatic burden at all four sites (Fig. 6B&C). Seven of the eight MPO inhibitor treated mice had no detectable bioluminescence metastases, with only one mouse developing lung metastasis, whereas all eight untreated 4T-1 BALB/c mice developed metastases across all four sites (Fig. 6D). Similar to the melanoma model, MPO inhibition decreased the frequencies of Ly6G⁺Ly6C⁺ myeloid cells across all metastatic sites, along with site-specific increases in CD8^+^ T cell and B cell frequencies compared to untreated 4T-1 BALB/c mice (Fig. 6E – G). In addition, MPO inhibition resulted in a significant reduction in Ly6G^+^ neutrophils across all metastatic sites (Fig. 6H) and site-specific changes in the frequencies of additional immune subsets resulting in reduced Ly6C⁺ monocytes and macrophages in the bone, as well as increased dendritic, NK, and CD4^+^ T cells (Fig. 6I – M). Phenotypic analysis of CD8^+^ T cells revealed an increase in the frequency of activated and effector CD8^+^ T cell subsets from MPO inhibition compared to untreated BALB/c mice but only within the lung tissues (Fig. S13A – F). Site specific differences in CD4^+^ T cell subsets were observed with an increase of CD44^+^, CD44^+^CD62L^+^ and CD25^+^ CD4^+^ T from MPO inhibition compared to untreated BALB/c mice within the bone and a decrease in activated CD4^+^ T in the lungs (Fig. S13G – L). BALB/c mice also tolerated the MPO inhibitor treatment well, with no significant changes in body weight (Fig. S13M). Together, these findings extend the anti-metastatic efficacy of adjuvant MPO inhibition to breast cancer and reveal shared, yet tissue-specific, alterations in the immune landscape of metastatic tissues.

**Figure 6.**
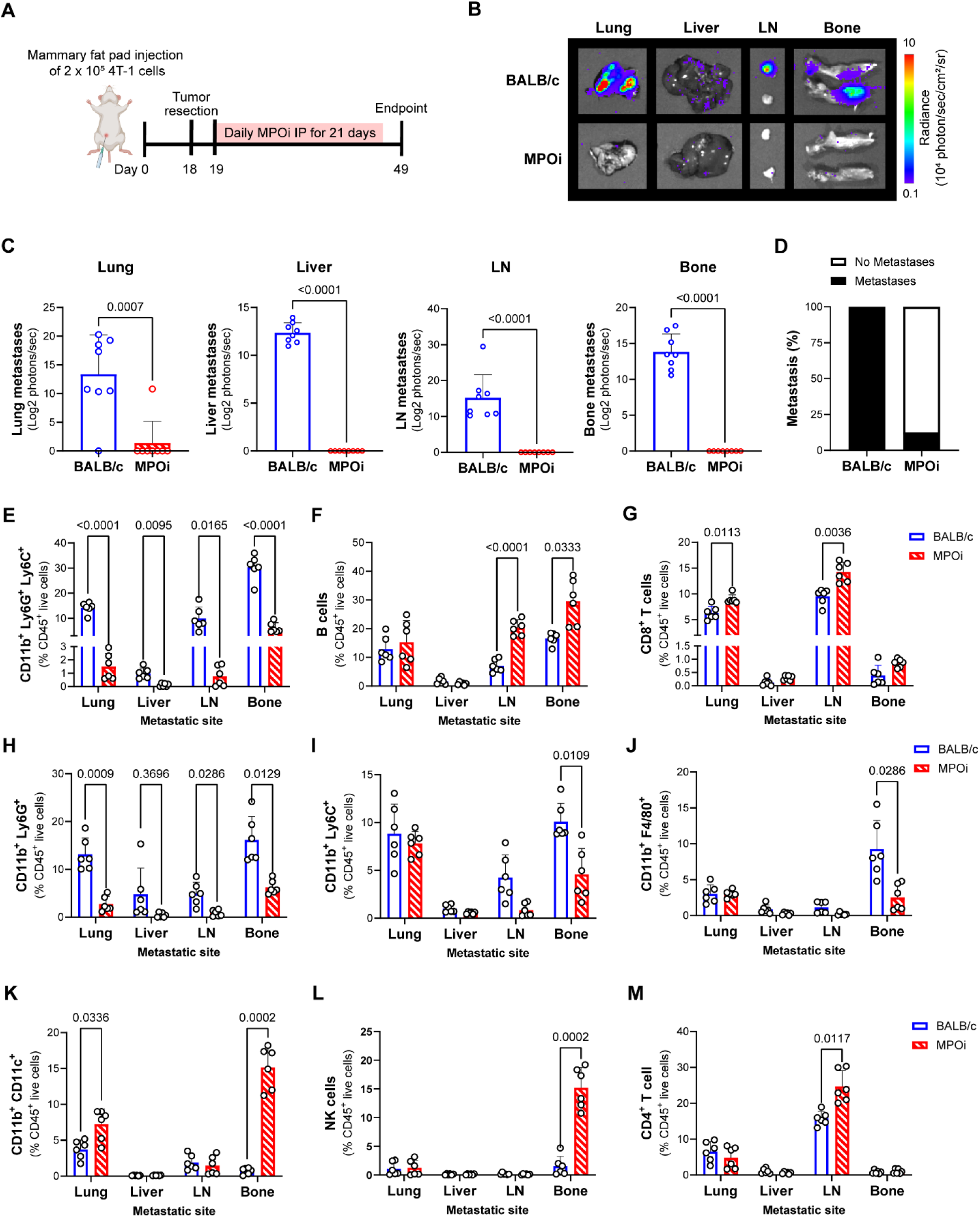
MPO inhibition significantly reduces multi-organ breast cancer metastatic burden. (A) Schematic of the spontaneous 4T-1 breast cancer metastasis model and MPO inhibitor, mitiperstat, (MPOi) treatment consisting of daily intraperitoneal injections for 21 days beginning on day 19. Representative (B) *ex vivo* B16F10 FLuc-eGFP bioluminescence imaging and corresponding (C) quantification of metastatic burden in the lung, liver, inguinal lymph nodes (LN) and bone, and (D) percentage of mice with detectable metastasis at experimental endpoint in BALB/c and MPOi mitiperstat treated BALB/c mice (n = 8 female mice/group). Flow cytometry quantification of frequencies (%) of (E) Ly6G^+^Ly6C^+^ myeloid cells, (F) B cells, (G) CD8^+^ T cells, (H) CD11b^+^Ly6G^+^ neutrophils, (I) CD11b^+^Ly6C^+^ monocytes, (J) CD11b^+^F4/80^+^ macrophages, (K) CD11b^+^CD11c^+^ dendritic cells, (L) NK cells and (M) CD4^+^ T cells in metastatic tissues from spontaneous 4T-1 metastatic BALB/c and MPOi treated BALB/c mice at experimental endpoint (n = 6 female mice/group). Data shown as mean ± SD; statistical significance of differences calculated using a two-way ANOVA followed by Tukey’s multiple comparisons test or Šídák’s multiple comparisons test (p values shown on graphs).

## Discussion

Immunosuppressive myeloid cells have been shown to promote metastasis, however, there are limited known therapeutically targetable pathways to inhibit their pro-metastatic functions. Here, we identify MPO as a druggable regulator of metastatic progression using a spontaneous metastatic melanoma model. We demonstrate that genetic MPO deficiency and adjuvant MPO inhibition markedly reduced multi-organ metastatic burden, accompanied by a reduction in immunosuppressive Ly6G^+^Ly6C^+^ myeloid cells and enrichment of B and T cell populations within metastatic tissues. Notably, the anti-metastatic efficacy of MPO inhibition was dependent upon B cells, CD8^+^ T cells and CD4^+^ T cells, as depletion of each population independently reversed MPO inhibitor-mediated metastatic control. Adjuvant MPO inhibition rapidly reduced circulating and disseminated tumor cells. We further validated these findings in a spontaneous breast cancer metastasis model using a distinct mouse genetic background, in which adjuvant MPO inhibition similarly reduced of multi-organ metastatic burden with comparable changes in the metastatic immune microenvironment. Together, these findings identify MPO as a therapeutically actionable regulator of metastatic progression and suggest that targeting MPO may limit metastatic disease by disrupting immunosuppressive myeloid cells.

While NOX2 was previously shown to contribute to experimental B16F10 lung metastasis^13^, NOX2-deficiency failed to reproduce the broad multi-organ metastatic reduction from MPO targeting observed in our spontaneous B16F10 metastasis model, suggesting that reduced NOX2-dependent ROS production alone was insufficient to explain the effects of MPO inhibition. Instead, MPO inhibition markedly reduced NET release, while NET deficiency and inhibition recapitulated the anti-metastatic effects of targeting MPO, implicating NETs as a potential downstream mediator of MPO-driven metastasis. While it is well established that NETs promote metastasis^44^, clinical strategies targeting NETs have had limited success. Our findings suggest that MPO inhibitors may represent an alternative therapeutic approach to inhibit NETs and subsequent metastatic disease. However, MPO may promote metastatic progression through additional mechanisms beyond NETs. We found that MPO deficiency and inhibition significantly reduced IL-1α secretion by Gr-1+ myeloid cells. IL-1α is a proinflammatory cytokine implicated in tumor progression and has been shown to found to inhibit T and NK cell activation and suppress the killing capacity of CD8^+^ T cells as well as recruit immunosuppressive myeloid populations such as MDSCs and tumor-associated macrophages.^45,46^ Future studies will evaluate whether MPO-dependent IL-1α signaling contributes to metastatic disease. Importantly, several MPO specific inhibitors have advanced to clinical trials for neurodegenerative and cardiovascular disorders, providing human safety, tolerability and pharmacokinetics data.^38,39,47–52^ Although our studies focused on mitiperstat, these clinically evaluated MPO inhibitors may offer opportunities for therapeutic repurposing in metastatic disease.

Our findings reinforce the established role of T cells in metastatic control and demonstrate that MPO contributes to myeloid cell-mediated suppression of T cell proliferation and activation. Previous studies have also demonstrated that B cells have anti-tumor properties where their depletion results in increased lung metastasis.^53,54^ Here, we extend those observations by demonstrating that B cells were required for MPO inhibitor-mediated metastatic control. Interestingly, MPO deficiency or inhibition attenuated myeloid cell-mediated suppression of B cell proliferation while reducing the secretion of IL-10 and IgM. Given the established role of IL-10 producing regulatory B cells in suppressing anti-tumor T cell responses^55^, these findings raise the possibility that targeting MPO may reduce regulatory B cell activity. Moreover, the reduction in IgM levels suggests that MPO inhibition may also shift B cells towards non-antibody mediated functions. Future studies will determine the mechanism by which MPO in myeloid cells influences B cell phenotypes and function during metastatic progression.

Although MPO inhibition was well tolerated in our preclinical models, the optimal dosing, treatment duration and long-term safety remain to be established. Notably, MPO deficiency is well tolerated in humans, with over 95% of affected individuals remaining asymptomatic^21,22,56^, supporting the potential clinical tolerability of MPO inhibition. Additionally, since MPO functions downstream of NOX2, MPO inhibitors are expected to preserve NOX2-dependent ROS generation, potentially maintaining protection against pathogens. While we have previously demonstrated that MPO inhibition enhances immune checkpoint inhibitor efficacy in subcutaneous tumor models including melanoma^19,34^, whether MPO inhibition similarly improves immunotherapy in the metastatic setting remains to be studied. Although genetic and pharmacologic disruption of NET formation reproduced the anti-metastatic effects of MPO targeting and several similar alterations in the metastatic immune microenvironment, the precise contribution of NETs to MPO-mediated metastatic progression remains to be identified, particularly given the effects of HCQ beyond NET inhibition. While our findings demonstrate that adjuvant MPO inhibition reduced circulating and disseminated tumor cells and metastatic outgrowth, its relative effects on each stage of the metastatic cascade remain to be resolved. Although MPO inhibition similarly reduced metastatic burden and altered the metastatic immune microenvironment in a spontaneous 4T-1 breast cancer model, future studies will define the underlying mechanisms and identify the immune populations responsible for MPO inhibitor-mediated metastatic control. Finally, our studies only evaluated adjuvant MPO inhibition and future studies will evaluate whether neoadjuvant MPO inhibitor treatment further prevents metastatic progression given the early immune remodeling observed from targeting MPO. Our findings demonstrate that MPO regulates Gr-1+ myeloid-mediated immunosuppression where depletion of Ly6G^+^ myeloid cells was not as effective as MPO inhibition at reducing metastasis suggesting that the contribution of MPO in supporting metastatic progression may extend beyond Ly6G⁺ subsets. Thus, the contributions of MPO within distinct myeloid populations require further investigation. Overall, our findings identify MPO as a therapeutically actionable regulator of metastatic progression and support MPO inhibitors as a promising strategy to control metastatic disease by inhibiting myeloid-mediated suppression and enhancing B and T cell mediated metastatic control.

## Methods

### MPO gene expression analysis

Gene expression data for skin cutaneous melanoma were obtained from The Cancer Genome Atlas (TCGA) via the UCSC Xena Browser (https://xenabrowser.net). The TCGA Skin Cutaneous Melanoma (TCGA-SKCM) cohort was analyzed using RNA sequencing data generated on the Illumina HiSeq platform and processed through the UCSC Xena pipeline. Gene expression values were provided as normalized counts and transformed as log2(normalized count + 1). Expression values for *MPO* (myeloperoxidase) were extracted based on gene symbol annotation. Samples were annotated according to TCGA sample type codes and categorized as primary tumor or metastatic tumor. Differences in *MPO* expression between primary and metastatic tumor samples were assessed using an unpaired t-test. Statistical significance was defined as *p* < 0.05.

### Reagents

Luminol (sodium salt), and lucigenin (N,N′-Dimethyl-9,9′-biacridinium dinitrate) were purchased from Sigma-Aldrich (MO, USA). L-012 sodium salt was purchased from Wako Chemicals USA (Richmond, VA, USA). Anti-CD20 (clone MB20-11), anti-CD8α (clone 2.43), anti-CD4 (clone GK1.5), anti-NK1.1 (clone PK136), anti-Ly6G (clone 1A8), and polyclonal rat IgG (clone C1.18.4) control were obtained from Bio X cell (BioXCell, NH, USA). Mitiperstat was purchased from MedChemExpress (MedChemExpress LLC, NJ, USA). Hydroxypropyl-β-cyclodextrin (HP-β-CD) was purchased from Thermo Fisher Scientific (Thermo Fisher Scientific, MA, USA). Luminol sodium salt was dissolved in sterile phosphate-buffered saline (PBS) to a final concentration of 50 mg/mL or 100 mg/mL and stored at -20°C. L-012 powder was dissolved in sterile double distilled water (ddH_2_0) to a final concentration of 20 mM and stored at -20°C. Mitiperstat was dissolved in DMSO to a final concentration of 358.42mM stored at −20°C. Hydroxychloroquine was dissolved in autoclaved water at a concentration of 0.5 mg/mL in cage bottles where cage bottles were replaced weekly.

### Cells

B16F10 cells stably expressing a ubiquitin C promoter driven click beetle green luciferase and enhanced green fluorescent protein (B16F10 CBG-GFP) were previously described.^35,36^ 4T-1 cells stably expressing a firefly luciferase and GFP (4T-1 FLuc-GFP) were kindly provided by Dr. Michale Bachmann from Michigan State University.

*In vivo spontaneous metastasis model.* The Institutional Animal Care and Use Committee at West Virginia University approved all animal protocols (2109047227). For the spontaneous melanoma metastasis model, 5 x 10^4^ B16F10 CBG-GFP cells were injected subcutaneously on the lower back of male and female 8-week-old C57BL/6 (The Jackson Laboratory, ME, USA) or age matched syngeneic C57BL/6 myeloperoxidase-deficient *MPO^-/-^* (MPO^tm1Lus^, The Jackson Laboratory, ME, USA) animals. Prior to tumor cell injection, the fur was removed on the lower back. Tumors were measured by calipers every 3 – 4 days once palpable. At 18 days post injection, primary subcutaneous tumor was surgically resected. For the breast cancer spontaneous metastasis model, 2 x 10^5^ 4T-1 cells were injected into the left 4th mammary fat pad of 8-week-old female BALB/c mice (The Jackson Laboratory, ME, USA). At 18 days post injection, primary mammary tumors were surgically resected. Mice were monitored twice weekly and weights measured until endpoint between day 39 – 42 post tumor injection for B16F10 and day 49 for 4T-1 cells. *In vivo* bioluminescence imaging of metastatic spread was performed 9 minutes after an intraperitoneal (IP) injection of D-luciferin (150 mg/kg) using an IVIS spectrum (Revvity, MA, USA) using the following acquisition parameters: acquisition time, auto; binning, 8; FOV, 13.2; f/stop, 1; filter, open. *In vivo* MPO activity and ROS levels were assessed using IP injections of luminol (200 mg/kg of body weight) and L-012 bioluminescence imaging (25 mg/kg of body weight). Images were acquired 9 min post each bioluminescence substrate injection (acquisition time, auto exposure up to 5 min; binning, 8; FOV, 6.6; f/stop, 1; filter, open). Luminol imaging occurred prior to L-012 imaging. At endpoint, metastatic burden was quantified using *ex vivo* bioluminescence imaging following IP injection of D-luciferin (150 mg/kg). At 5 minutes post luciferin injection, mice were euthanized, and metastatic tissues were immediately excised and imaged on an IVIS spectrum using the same acquisition parameters described above. Bioluminescence data were analyzed in Living Image software (Revvity, MA, USA). A background region of interest (ROI) was drawn on every image to measure background bioluminescence. Luciferase, luminol and L-012 bioluminescence was quantified by drawing a uniform background-subtracted ROI around the torso of the mouse for *in vivo* imaging or around each excised tissue for ex vivo imaging. Using a background-subtracted ROI to measure the bioluminescence accounts for any fluctuations in noise in every image where signal output is the measured reporter’s photon flux above background. Total photon flux for each mouse was exported to Microsoft Excel and graphed using GraphPad Prism (GraphPad Software Inc., CA, USA) where total photon flux (Y) was transformed using the following equation: Log2(Y + 1). For MPO inhibition studies, mice received daily intraperitoneal (IP) injections of 200 µL mitiperstat (180 µmol/kg in 5% DMSO, 95% of 15% HP-β-CD [w/v] in PBS) for 21 days beginning one day post tumor resection. For the immune cell depletion studies, antibodies against CD8, CD4, NK1.1, Ly6G or IgG were injected into animals (250 µg in 100 µL PBS intraperitoneally) twice weekly for 3 weeks beginning on day 19 post tumor inoculation. A single intraperitoneal injection of anti-CD20 (250 µg in 100 µL PBS) occurred for B cell depletion on day 19 post tumor inoculation. An IgG isotype antibody was used as a control. For enumerating circulating and disseminated melanoma cells, blood was collected and red blood cells were lysed and single cell suspensions were harvested from metastatic target tissues of the lungs, lymph nodes and bone marrow. B16F10 GFP positive cells were quantified on Cytek® Aurora (Cytek, MD, USA).

### Cresyl violet staining

Metastatic tissues at experimental endpoint were fixed in 10% neutral buffered formalin, embedded in paraffin, and sectioned at a thickness of 4–6 µm using a microtome. Slides were deparaffinized for 20 minutes with HistoClear II (Electron Microscopy Sciences, PA, USA), then cleared with ethanol gradients for 3 minutes each, 100%, then 90%, then 70%. They were then washed in cold PBS. Slides were immersed in 0.01% Cresyl violet acetate solution (Abcam, MA, USA) for 5 minutes. After 5 minutes, slides were placed in distilled water to stop the staining, then placed in 100% ethanol for 2 minutes for stain differentiation. Slides were allowed to air dry before a coverslip was mounted and left to dry overnight. Slides were imaged on an Olympus MVX10 (Evident Scientific, MA, USA) 1x brightfield. Cresyl violet staining was quantified with ImageJ^57^ by drawing an ROI around the total tissue area and stained are to get percent of staining.

### Flow Cytometry

Single cell suspensions were harvested from metastatic tissues. Single cell suspensions were harvested from the spleen of age-matched healthy (tumor-free) mice. Tumors were dissociated using a mouse tumor dissociation kit (Miltenyi Biotec, CA, USA). Red blood cells are lysed using 1X red blood cell lysis buffer (Thermo Fisher Scientific, CA, USA). Remaining cells were resuspended in cell staining buffer (BioLegend, CA, USA) at a concentration of 2 x 10^5^ – 1 x 10^6^ cells per 100 µl. Fc receptors are blocked using 10 µg/ml ChromPure of mouse IgG (Jackson ImmunoResearch Inc., PA, USA) per 10^6^ cells in a 100 µl volume for 10 min on ice. Cells were washed with cell staining buffer and incubated with an antibody mix to label immune and T cells (Table S1 & S2) for 15 – 20 minutes on ice in the dark. Cells were fixed using 200 µl Fixation buffer (Thermo Fisher Scientific, CA, USA) for 20 min at 4°C in the dark. Cells were washed 2 times with 1X PermWash (BD Bioscience, CA, USA) and permeabilized using Permeabilization solution (BD Bioscience, CA, USA). Intracellular antibodies were incubated for 20 min at room temperature in the dark. Stained cells were analyzed using the Cytek® Aurora (Cytek, MD, USA) within 2 weeks of staining. Data was analyzed using FCS express (De Novo Software, CA, USA). Gating strategies for immune cells, CD8^+^ and CD4^+^ T cells are shown in figures S14 - 16.

*Gr-1+ myeloid cells cell isolation:* At tumor endpoint, single cell suspensions were harvested from the metastatic sites of spontaneous metastatic mice. Single cell suspensions were harvested from the spleen of age-matched healthy (tumor-free) mice. Cd11b^+^Gr-1^+^ myeloid cells were isolated using an EasySep™ mouse MDSC isolation kit (STEMCELL Technologies Inc, MA, USA). Cd11b^+^Gr-1^+^ myeloid cells were quantified by flow cytometry using CD11b-APC (Miltenyi Biotec, CA, USA) and Gr-1-PE (Miltenyi Biotec, CA, USA) and only Cd11b^+^Gr-1^+^ myeloid cells purity of >80% was used in the subsequent studies quantifying T cell proliferation suppression, reactive oxygen species levels and extracellular trap formation. *MPO activity and ROS quantification*.

MPO activity was assessed by luminol and ROS were assessed using L-012 of the isolated Gr-1+ myeloid cells. 1 x 10^5^ isolated Gr-1+ myeloid cells were added to 100 mM luminol or 50 μM L-012 in colorless DMEM media in a 96-well black walled plate. Cells were immediately imaged for 60 min using the Kino imaging system (Spectral Instruments Imaging, AZ, USA) at 37°C under 5% CO_2_ flow. Typical acquisition parameters were as follows: acquisition time, 5 min; binning, 16; FOV, 13 cm; f/stop, 1.2; filter, open; total number of acquisitions 12. Bioluminescence photon flux (photons/sec) data were analyzed by ROI measurements with background subtraction in Aura (Spectral Instruments Imaging, AZ, USA); these raw data were imported into Excel (Microsoft Corp., WA, USA), averaged in each individual experiment done in duplicate or triplicate wells. The area under the curve was calculated from images taken at 5 – 60 min. The area under the curve for each reporter was taken from n = 6 independent isolation experiments.

### NET quantification

1 x 10^5^ isolated Gr-1+ myeloid cells in 100 μL RPMI 1640 media were incubated for 1 – 2 hours in a 96 well plate. After incubation, the cells were spun at 1000 rpm for 10 min. 50 μL of supernatant was removed and mixed with 50 μL of 1X TE buffer (Thermo Fisher Scientific, CA, USA) in a 96 well black walled clear bottom plate. 50 μL of 1:200 dilution of Pico Green (Thermo Fisher Scientific, CA, USA) in 1X TE buffer was added to each well and fluorescence (Excitation: 480nm and Emission: 520nm) was quantified on a Synergy HTX multi-mode plate reader (BioTex, CA, US). A standard curve was made using the lambda DNA standard (Thermo Fisher Scientific, CA, USA) in 1X TE buffer (concentration range from 1 μg/ml to 25 pg/ml). The DNA concentration of the samples were determined from the generated DNA standard curve.

### T and B cell proliferation assay

CD8^+^ T cells and CD19^+^ B cells were isolated from the spleen of 8 – 10-week-old, sex-matched healthy C57BL/6 mice using an EasySep™ mouse CD8^+^ T cell isolation kit or EasySep™ Mouse B Cell Isolation Kit (STEMCELL Technologies Inc, MA, USA). 1 x 10^6^ isolated T or B cells/mL were stained with 5 μM concentration of CellTrace CFSE (Thermo Fisher Scientific, CA, USA) and incubated at 37°C for 20 min. Cells were incubated for 5 min with culture media to remove unbound dye and subsequently washed 2 times. CSFE stained T or B cells were incubated for 2 – 3 hours at 37°C in a humidified 5% CO_2_ atmosphere at a final concentration of 1 x 10^5^ isolated cells/ 100 μL. T cells were stimulated using a 1:1 bead-to-cell ratio of mouse T cell activation CD3/CD28 dynabeads (5 μL, Thermo Fisher Scientific, CA, USA), while B cells were stimulated with 10ng/mL IL-4 (R&D Systems, MN, USA). Controls were fixed using 0.4% paraformaldehyde overnight after CSFE staining of T and B cells: 2 x 10^5^ unstained (no CSFE) T cells or B cells, 2 x 10^5^ CSFE stained T cells or B cells (time zero, t = 0), 2 x 10^5^ Gr-1^+^ myeloid cells. Using a 1:1 ratio, 2 x 10^5^ T cells or B Cells were cultured with 2 x 10^5^ Gr-1^+^ myeloid cells in 500 μL of RPMI Medium 1640 with 2 mM L-Glutamine, 10% FBS, and 100 U/ml penicillin/streptomycin and 5 μL dynabeads, for T cells, or 10ng/mL IL-4 (R&D Systems, MN, USA), for B cells, in 24-well plates for 72 hours at 37°C in a humidified 5% CO_2_ atmosphere. After 72 hours, the 24-well plate was washed 2 times with culture media and the cell pellets were fixed using 0.4% paraformaldehyde overnight in the dark. T cell proliferation was quantified using the BD LSRFortessa™ Cell Analyzer (BD Biosciences, NJ, USA). Data was analyzed using FCS express (De Novo Software, CA, USA).

#### ELISA

Culture supernatants collected from the T and B cell co-cultures with Gr-1+ myeloid cells were stored at –20°C for subsequent analysis using a mouse TNFα, IL-10, IgM, IgA, IgG ELISA kit (Invitrogen, CA, USA) following the manufacturer’s protocol. Inactivated T and B cell alone controls were included. A standard curve was generated using the standards provided in the kit, and sample concentrations were calculated by interpolation from the standard curve using Microsoft Excel.

*Gr-1+ myeloid cell cytokine measurement.* A total of 1.5 x 10^5^ Gr-1+ myeloid cells in 150 μl of RPMI media containing 100 U/ml penicillin/streptomycin were seeded in 96-well plates and incubated at 37°C in 5% CO2. After a 24-hour incubation, supernatants were collected, centrifuged at 350 rpm for 5 minutes at 4°C, and stored at –80°C until further use. Cytokine levels were measured using a mouse 48-target Olink cytokine assay (Genewiz, NJ, USA).

### Transwell migration studies

Bone marrow was isolated from the hind limbs of healthy 10-week old male and female C57BL/6 mice and filtered through a 70µm cell strainer (Fisher Scientific, Waltham, MA). Single cells were counted and resuspended at a concentration of 2.5 x 10^6^ cells/mL in RPMI 1640 media supplemented with 20 ng/mL GM-CSF, 10% FBS, and 1X penicillin/streptomycin (Gr-1+ media). 5 x 10^6^ cells in 2 mL of Gr-1+ media were plated per well in a 6-well plate. On day 3, the number of non-adherent cells in the supernatant were quantified, where Gr-1+ cells were then isolated from these non-adherent cells using the EasySep™ mouse MDSC isolation kit (STEMCELL Technologies Inc, MA, USA) according to the manufacturer’s instructions. Gr-1+ myeloid cells were incubated with Gr-1+ media (WT), 50 µM Mitiperstat or 50 µM HCQ for 20 min. To assess migration of B16F10 cells toward Gr-1+ myeloid cells, a transwell assay (Falcon, Corning, NY) was performed with 1 x 10^5^ B16F10 CBG-GFP cells in 500 µL of B16F10 cell media seeded in the upper chamber and 1 x 10^5^ Gr-1+ myeloid cells in 500 µL of MDSC media in the lower chamber. Migration of B16F10 cells toward Gr-1+ myeloid cells was quantified by enumerating the % of GFP-positive tumor cells from the supernatant and from trypsinized adherent cells detached from the lower well. To assess migration of Gr-1+ myeloid cells toward B16F10 cells, Gr-1+ myeloid cells were labeled with 5 µM CFSE for 20 minutes prior to seeding in the upper chamber and parental B16F10 cells in the lower chamber using the same densities described above. Migration of Gr-1+ myeloid cells to B16F10 was quantified by enumerating the % of CSFE-positive Gr-1+ myeloid cells from the supernatant.

### In vivo co-injection spontaneous metastasis model

5 × 10^4^ Gr-1+ myeloid cells isolated from the spleen of metastasis-bearing WT, MPO inhibitor treated or HCQ treated mice were mixed in a 1:1 ratio with B16F10 CBG-GFP cells and subcutaneously injected into the right hind limb of recipient WT mice. Primary subcutaneous tumor resection and subsequent metastasis was monitored as described above in the in vivo spontaneous metastasis model section.

### Statistical analyses

Graphs were made and statistical analyses were performed using GraphPad Prism (GraphPad Software, Inc, CA, USA). Data were expressed as mean ± SD. For analysis of three or more groups, analysis of variance (ANOVA) tests were performed followed by a Tukey’s or Bonferroni’s multiple comparisons test, or false discovery rate test. Standard comparison of survival curves used the log rank (Mantel-Cox) analysis. Analysis of differences between two normally distributed paired test groups were performed using a Student’s t-test. P values were considered statistically significant if *p* < 0.05.

### Graphical Illustrations

Graphical elements used to create experimental design schemes were created in Bio Render licensed by West Virigina University Department of Microbiology, Immunology and Cell Biology.

## Acknowledgements

Imaging experiments and image analysis were performed in the West Virginia University (WVU) Animal Models & Imaging Facility which has been supported by the WVU Cancer Institute, the WVU HSC Office of Research and Graduate Education, and NIH grants P20GM121322 and U54GM104942. Flow cytometry studies were performed in the West Virginia University Flow Cytometry & Single Cell Core Facility, RRID:SCR_017738 which has been supported by TME CoBRE GM121322, S10 equipment grant OD028605, WVCTS grant #GM104942 and WV-INBRE grant #GM103434. Specimens were provided by the West Virginia University Biospecimen Translational Research Analysis Core (BIOTRAC) which has been supported by the WVU Cancer Institute and NIH grants P20GM103434 and U54GM104942.

