## Supplemental Information for "Adjuvant myeloperoxidase inhibition promotes B and T cell mediated control of murine melanoma metastasis"

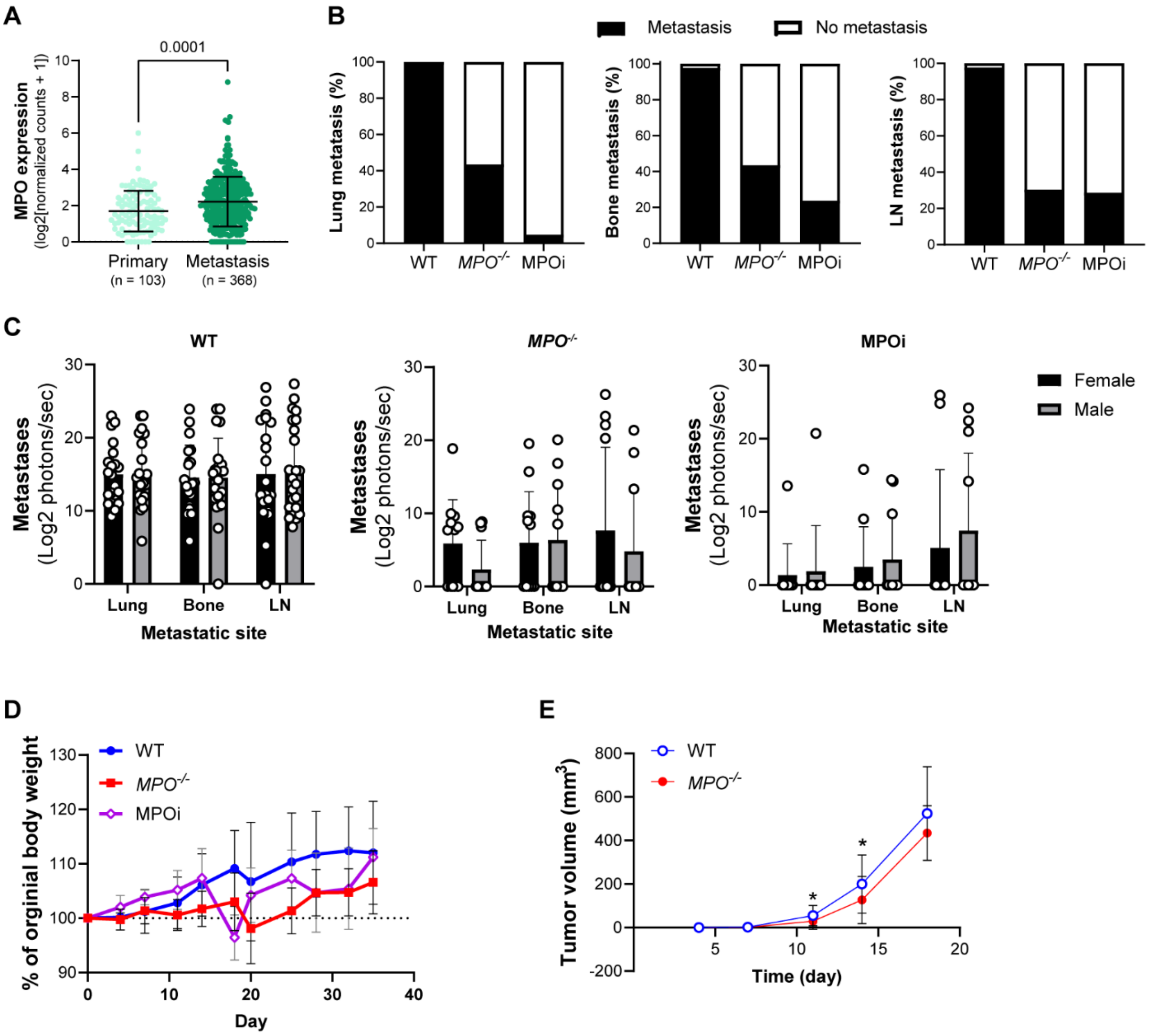


**Figure S1. MPO expression is increased in metastatic melanoma.** (A) TCGA melanoma (SKCM) analysis shows higher MPO mRNA transcript levels in metastasis (n = 368) compared to primary tumors (n = 103); data shown as mean ± SD where statistical significance of differences calculated using unpaired t test (p values shown on graphs). (B) Percentage of mice with detectable metastasis and (C) *ex vivo* bioluminescence quantification of metastatic burden comparing male and female mice in lung, bone and inguinal lymph nodes (LN) at experimental endpoint in WT, *MPO*$^{-/-}$ and MPO inhibitor, mitiperstat, (MPOi) treated WT spontaneous B16F10 metastatic mice (n = 44 WT, n = 23 *MPO*$^{-/-}$, n = 21 MPOi; equal sex). (D) Body weight measurements of WT, *MPO*$^{-/-}$ and MPOi mice and (E) subcutaneous tumor volume growth curve comparing WT and *MPO*$^{-/-}$ mice prior to tumor resection. Data shown as mean ± SD; statistical significance of differences calculated using a two-way ANOVA followed by Šídák's multiple comparisons test or unpaired t test where * $p < 0.05$.

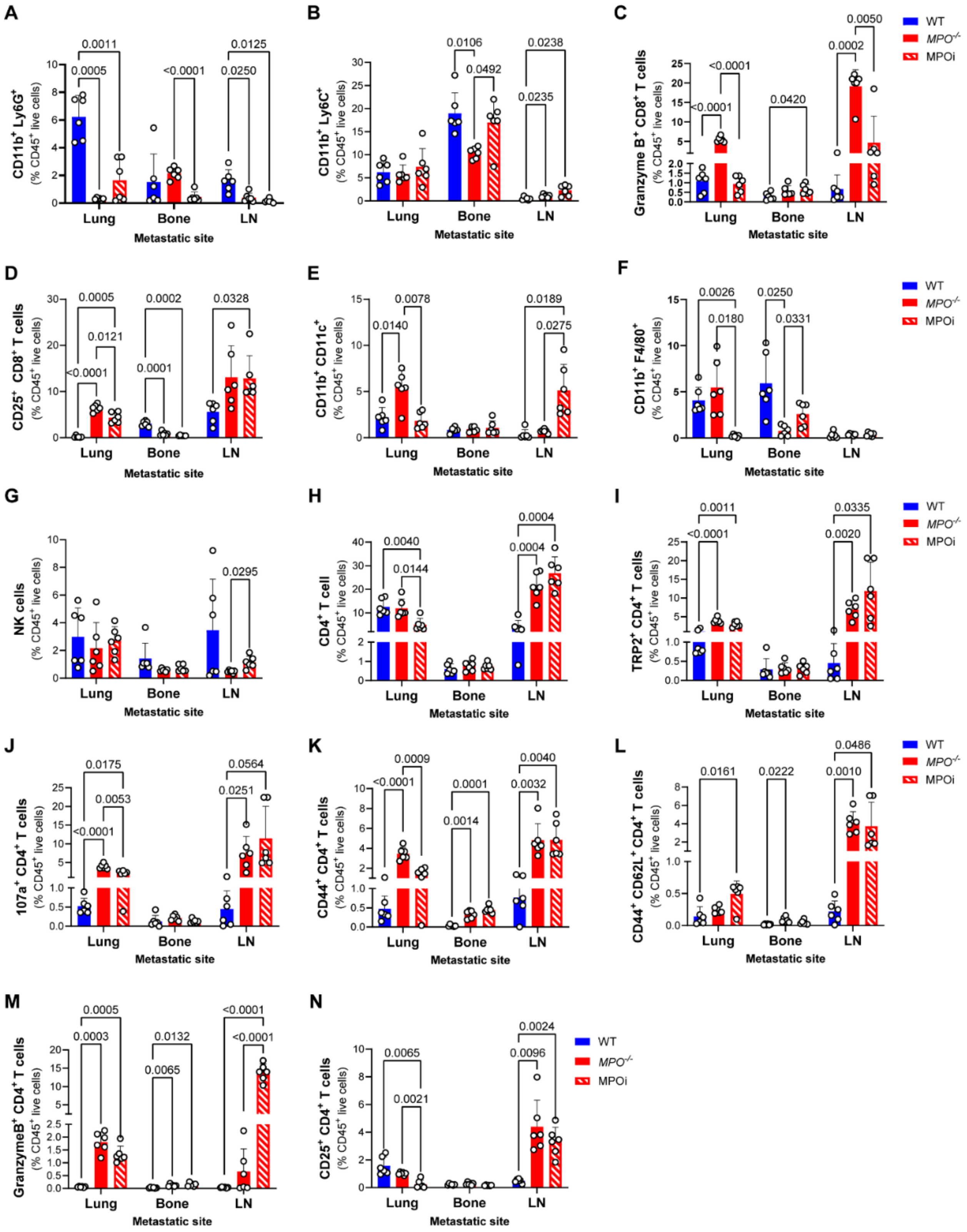


**Figure S2. MPO deficiency and inhibition remodels the metastatic immune microenvironment related to Figure 3.** Flow cytometry quantification of frequencies (%) of (A) $CD11b^+Ly6G^+$ neutrophils, (B) $CD11b^+Ly6C^+$ monocytes, (C) $GranzymeB^+$ $CD8^+$ T cells, (D)

$CD25^+$ $CD8^+$ T cells, (E) $CD11b^+CD11c^+$ dendritic cells, (F) $CD11b^+F4/80^+$ macrophages, (G) NK cells, (H) $CD4^+$ T cells, (I) $TRP2^+CD4^+$ T cells, (J) $CD107a^+CD4^+$ T cells, (K) $CD44^+CD4^+$ T cells, (L) $CD44^+CD62L^+CD4^+$ T cells, (M) $GranzymeB^+$ $CD4^+$ T cells, (N) D) $CD25^+$ $CD4^+$ T cells in the lungs, bone, and inguinal lymph nodes (LN) from spontaneous B16F10 metastatic WT, $MPO^{-/-}$ and MPOi mitiperstat treated WT mice at experimental endpoint. Data shown as mean ± SD (n = 6 mice/group, equal sex); statistical significance of differences in immune cell frequencies calculated using a two-way ANOVA followed by Tukey's multiple comparisons test (p values shown on graphs).

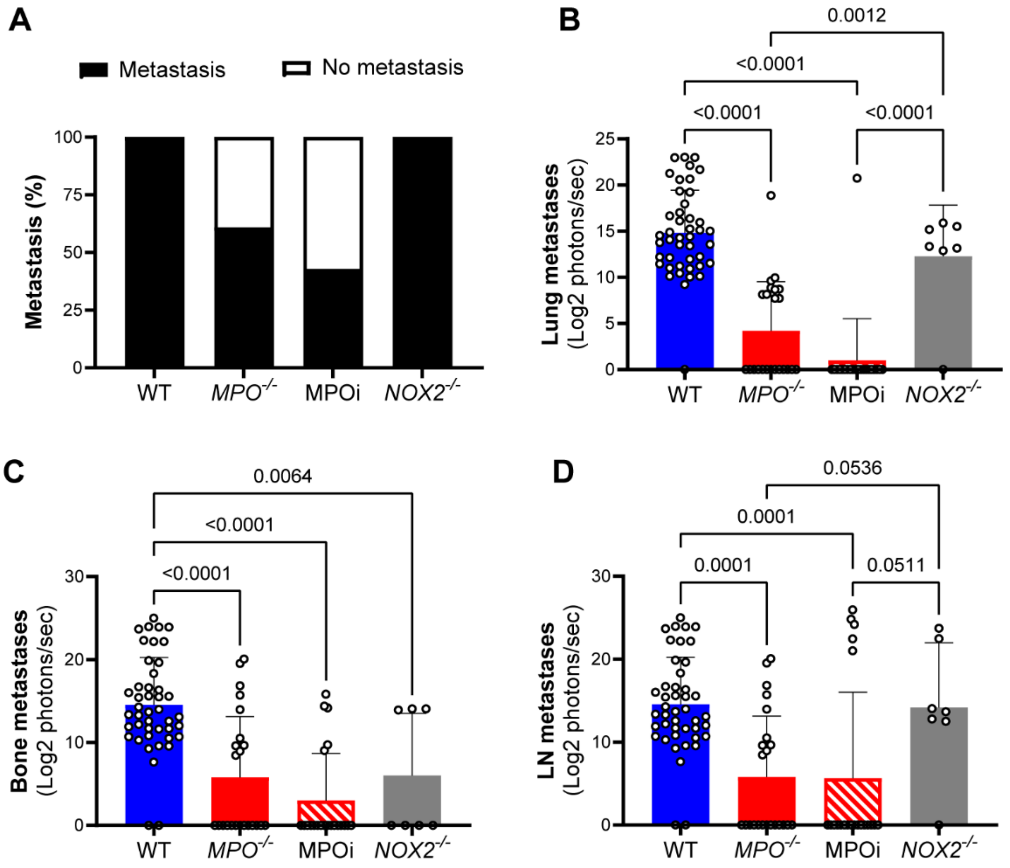


**Figure S3. NOX2 deficiency does not decrease metastatic melanoma burden as effectively as targeting MPO.** (A) Percentage of mice with detectable metastasis and *ex vivo* bioluminescence quantification of metastatic burden in (B) lung, (C) bone and (D) inguinal lymph nodes (LN) at experimental endpoint comparing NOX2-deficient (*NOX2*$^{-/-}$) and WT, *MPO*$^{-/-}$ and MPOi mitiperstat treated WT mice at experimental endpoint. Data shown as mean ± SD (n = 6 *NOX2*$^{-/-}$, equal sex, WT, *MPO*$^{-/-}$ and MPOi mice same dataset as shown in figure 2); statistical significance of differences in immune cell frequencies calculated using a two-way ANOVA followed by Tukey's multiple comparisons test (p values shown on graphs).

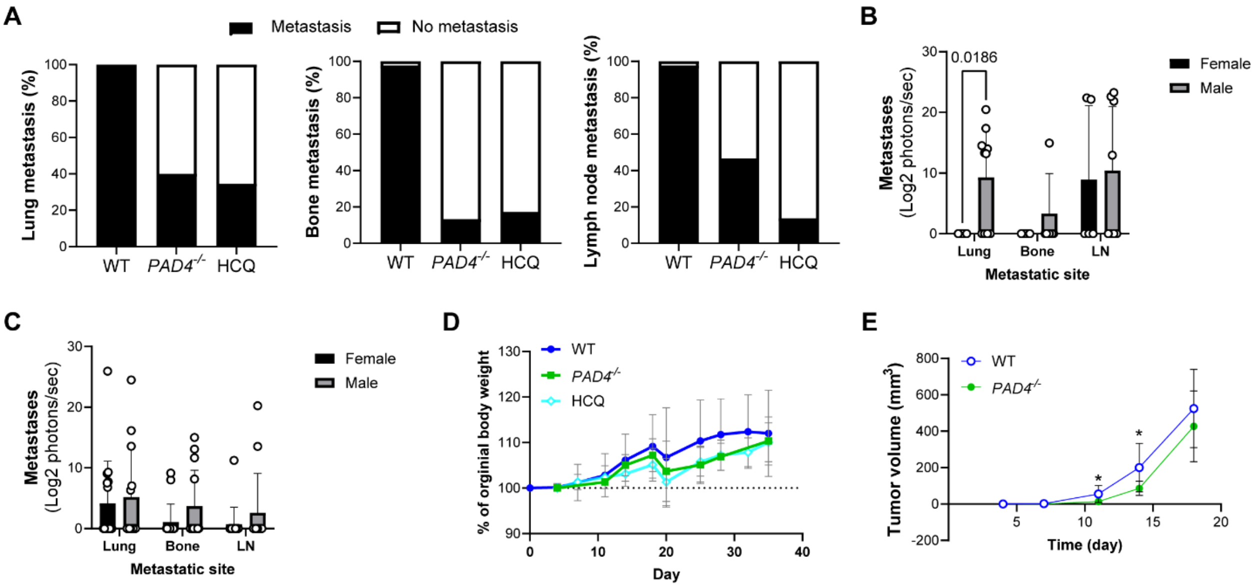


**Figure S4. NET deficiency and inhibition reduces melanoma metastasis related to Figure 4.** (A) Percentage of mice with detectable metastasis and *ex vivo* bioluminescence quantification of metastatic burden comparing male and female mice in lung, bone and inguinal lymph nodes (LN) at experimental endpoint in (B) *PAD4*$^{-/-}$ and (C) HCQ-treated WT spontaneous B16F10 metastatic mice. (D) Body weight measurements of WT, *PAD4*$^{-/-}$ and HCQ treated WT mice and (E) subcutaneous tumor volume growth curve comparing WT and *PAD4*$^{-/-}$ mice prior to tumor resection. Data shown as mean ± SD (n = 44 WT same dataset as shown in figure 2, n = 15 *PAD4*$^{-/-}$, n = 29 HCQ; equal sex); statistical significance of differences calculated using a two-way ANOVA followed by Šídák's multiple comparisons test or unpaired t test where p value is shown on graph or * $p < 0.05$.

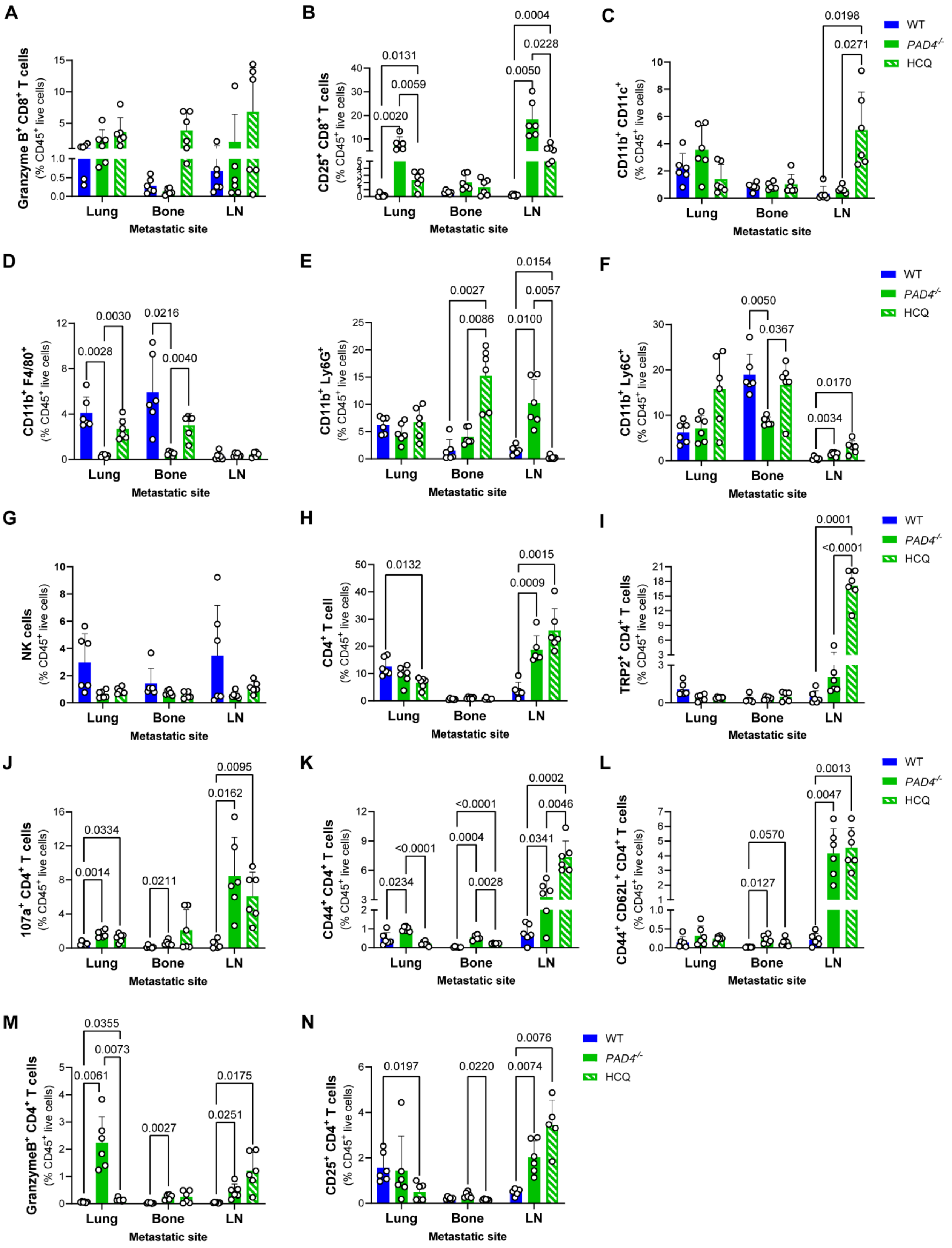

A
Granzyme B+ CD8+ T cells (% CD45+ live cells)
Lung
Bone
LN
Metastatic site
B
CD25+ CD8+ T cells (% CD45+ live cells)
0.0131
0.0059
0.0020
0.0004
0.0228
0.0050
Lung
Bone
LN
Metastatic site
C
CD11b+ CD11c+ (% CD45+ live cells)
0.0198
0.0271
Lung
Bone
LN
Metastatic site
WT
PAD4-/-
HCQ
D
CD11b+ F4/80+ (% CD45+ live cells)
0.0030
0.0028
0.0216
0.0040
Lung
Bone
LN
Metastatic site
E
CD11b+ Ly6G+ (% CD45+ live cells)
0.0154
0.0027
0.0057
0.0086
0.0100
Lung
Bone
LN
Metastatic site
F
CD11b+ Ly6C+ (% CD45+ live cells)
0.0050
0.0367
0.0170
0.0034
Lung
Bone
LN
Metastatic site
WT
PAD4-/-
HCQ
G
NK cells (% CD45+ live cells)
Lung
Bone
LN
Metastatic site
H
CD4+ T cell (% CD45+ live cells)
0.0132
0.0015
0.0009
Lung
Bone
LN
Metastatic site
I
TRP2+ CD4+ T cells (% CD45+ live cells)
0.0001
<0.0001
Lung
Bone
LN
Metastatic site
WT
PAD4-/-
HCQ
J
107a+ CD4+ T cells (% CD45+ live cells)
0.0334
0.0014
0.0211
0.0095
0.0162
Lung
Bone
LN
Metastatic site
K
CD44+ CD4+ T cells (% CD45+ live cells)
<0.0001
0.0234
<0.0001
0.0004
0.0028
0.0002
0.0046
0.0341
Lung
Bone
LN
Metastatic site
L
CD44+ CD62L+ CD4+ T cells (% CD45+ live cells)
0.0570
0.0127
0.0013
0.0047
Lung
Bone
LN
Metastatic site
WT
PAD4-/-
HCQ
M
GranzymeB+ CD4+ T cells (% CD45+ live cells)
0.0355
0.0073
0.0061
0.0027
0.0175
0.0251
Lung
Bone
LN
Metastatic site
N
CD25+ CD4+ T cells (% CD45+ live cells)
0.0197
0.0220
0.0076
0.0074
Lung
Bone
LN
Metastatic site
WT
PAD4-/-
HCQ

**Figure S5. NET deficiency and inhibition remodels the metastatic immune microenvironment related to Figure 4.** Flow cytometry quantification of frequencies (%) of (A) GranzymeB$^+$ CD8$^+$ T cells, (B) CD25$^+$ CD8$^+$ T cells, (C) CD11b$^+$CD11c$^+$ dendritic cells, (D) CD11b$^+$F4/80$^+$ macrophages, (E) CD11b$^+$Ly6G$^+$ neutrophils, (F) CD11b$^+$Ly6C$^+$ monocytes, (G) NK cells, (H) CD4$^+$ T cells, (I) TRP2$^+$CD4$^+$ T cells, (J) CD107a$^+$CD4$^+$ T cells, (K) CD44$^+$CD4$^+$ T cells, (L) CD44$^+$CD62L$^+$CD4$^+$ T cells, (M) GranzymeB$^+$ CD4$^+$ T cells, (N) D) CD25$^+$ CD4$^+$ T cells in the lungs, bone, and inguinal lymph nodes (LN) from spontaneous B16F10 metastatic WT, *PAD4*$^{-/-}$ and HCQ treated WT mice at experimental endpoint. Data shown as mean ± SD (n = 6 mice/group, equal sex; WT same dataset as shown in figure 3 and S2); statistical significance of differences in immune cell frequencies calculated using a two-way ANOVA followed by Tukey's multiple comparisons test (p values shown on graphs).

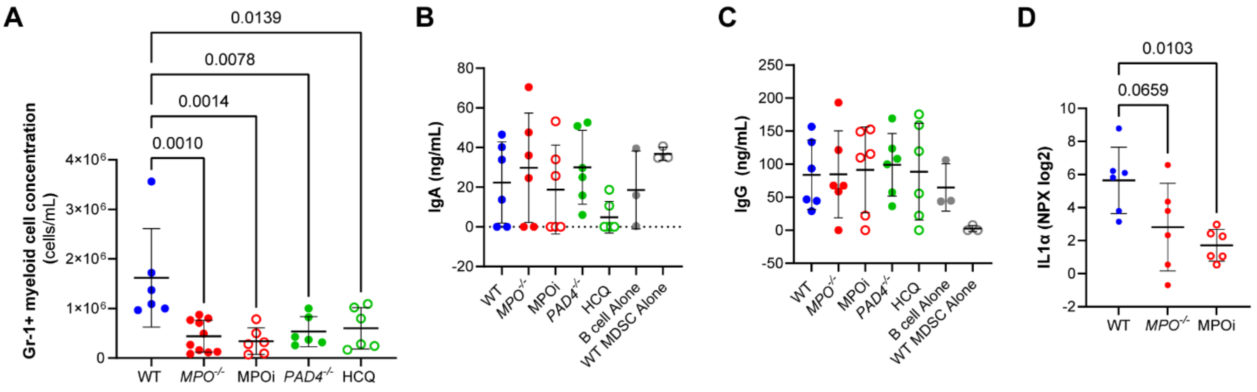


**Figure S6. Targeting MPO and NETs alters Gr-1+ myeloid cells.** Quantification of (A) lung-isolated Gr-1+ myeloid cell concentration, (B) IgA and (C) IgG levels in the supernatant of Gr-1+ myeloid and B cell co-culture studies and (D) IL-1α levels secreted by Gr-1+ myeloid cells isolated from the lung of spontaneous B16F10 metastatic mice at experimental endpoint in WT, *MPO*$^{-/-}$, MPOi, *PAD4*$^{-/-}$ and HCQ mice. Data shown as mean ± SD (n = 6 mice/group, equal sex); statistical significance of differences calculated using a two-way ANOVA followed by Tukey's multiple comparisons test (p values shown on graphs).

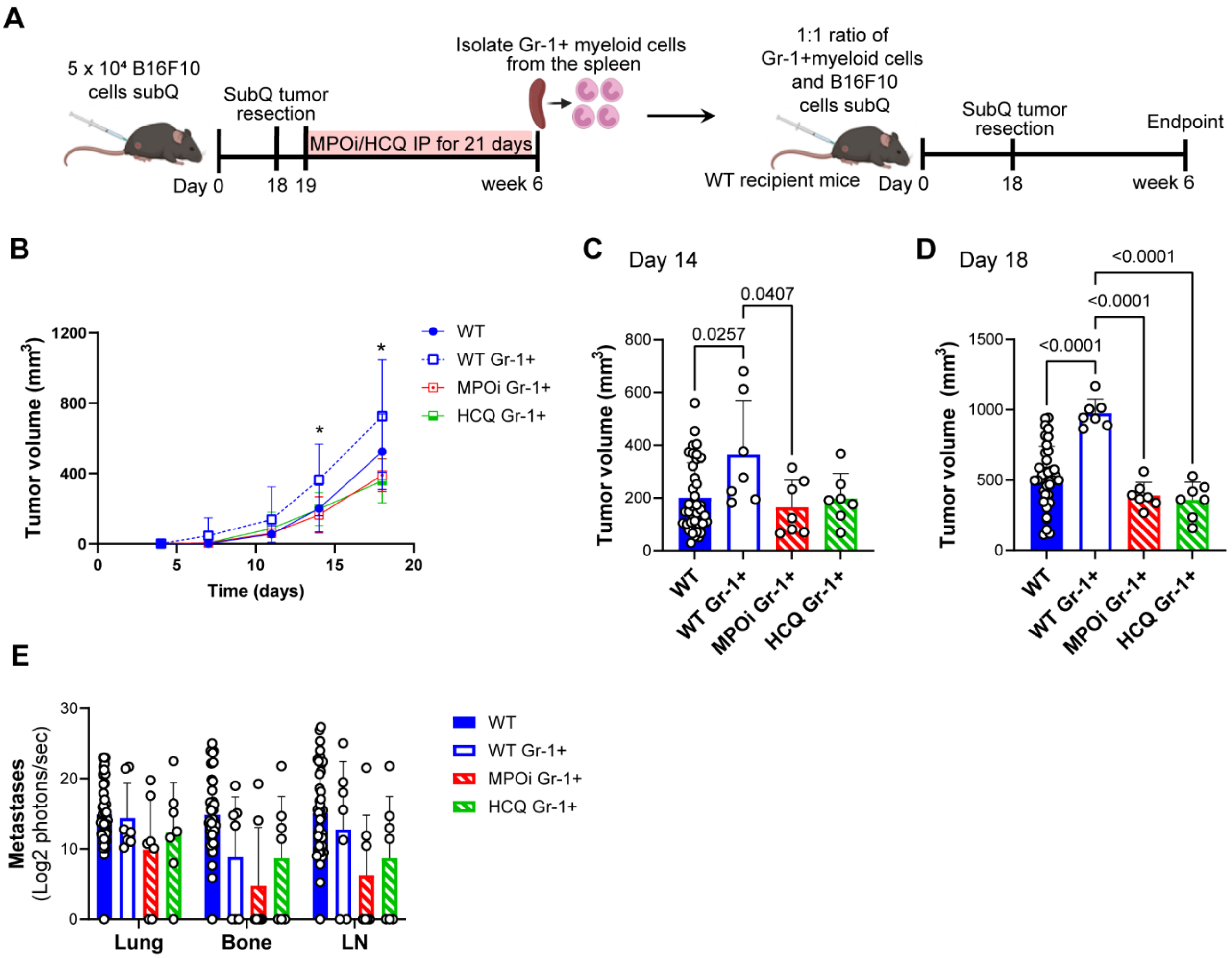


**Figure S7. MPO and NET inhibited Gr-1+ myeloid cells have reduced tumor-promoting abilities.** (A) Schematic of co-injection studies in the spontaneous B16F10 metastatic model where Gr-1+ myeloid cells in untreated WT, MPO inhibitor (MPOi) or HCQ treated WT mice were isolated from the spleen and injected in a 1:1 ratio with B16F10 CBG-GFP cells into recipient WT mice. (B) Tumor growth curve, tumor volume at (C) day 14 and (D) day 18, and (E) ex vivo bioluminescence quantification of metastatic burden at the experimental endpoint following co-injection of Gr-1+ myeloid and B16F10 cells. Untreated WT mice included for comparison (same dataset as in Figure 2). Data shown as mean ± SD (n = 7 mice/group, equal sex); statistical significance of differences calculated using a two-way ANOVA followed by Tukey's multiple comparisons test (p values shown on graphs or * $p < 0.05$).

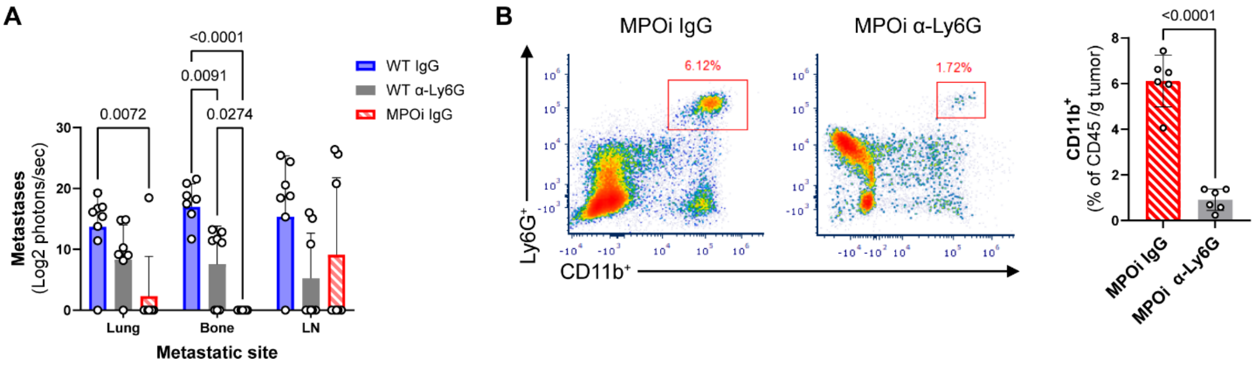


**Figure S8. Ly6G$^+$ myeloid cell depletion is not as effective as MPO inhibition.** (A) *Ex vivo* bioluminescence quantification of metastatic burden in lung, bone and inguinal lymph nodes (LN) at experimental endpoint following Ly6G$^+$ myeloid cell depletion in spontaneous B16F10 metastatic WT mice (α-Ly6G) compared to WT IgG treated (WT IgG) and MPO inhibitor IgG treated (MPOi IgG) WT mice at experimental endpoint. (B) Flow cytometry plots and corresponding quantification confirming depletion of Ly6G$^+$ myeloid cells from the spleen. Data shown as mean ± SD (n = 8 mice/group, equal sex, WT IgG and MPOi IgG mice same dataset as shown in figure 5); statistical significance of differences in immune cell frequencies calculated using a two-way ANOVA followed by Tukey's multiple comparisons test or unpaired t test (p values shown on graphs).

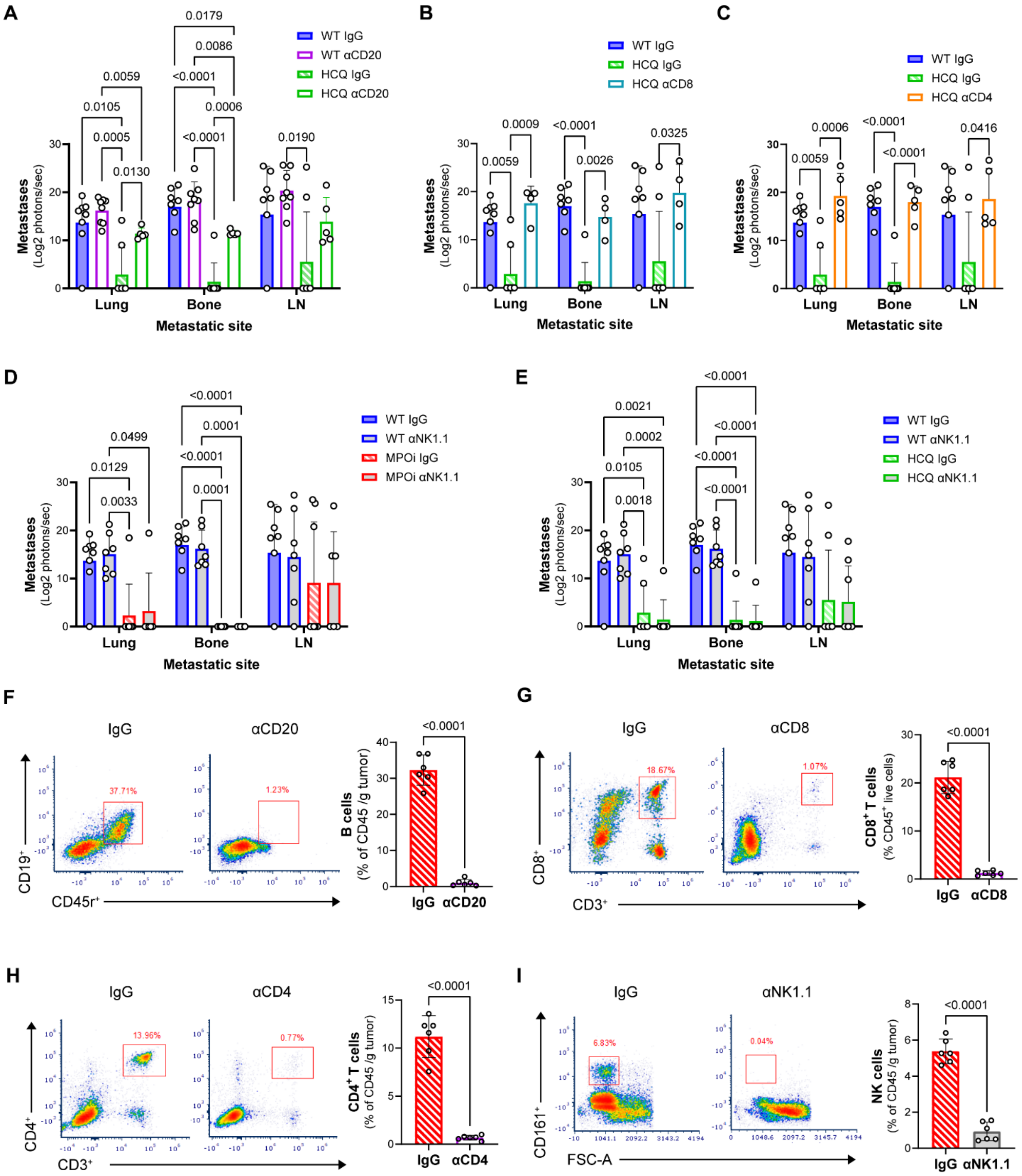

**Figure S9. Anti-metastatic effects of HCQ are dependent upon B, CD8⁺ and CD4⁺ T cells related to Figure 6.** *Ex vivo* bioluminescence quantification of metastatic burden in spontaneous B16F10 metastatic WT and HCQ treated mice in the lungs, bone and inguinal lymph nodes (LN) at experimental endpoint following selective depletion of (A) B cells using an anti-CD20 (αCD20) antibody, (B) CD8⁺ T cells using an anti-CD8α antibody (αCD8), and (C) CD4⁺ T cells using an anti-CD4 antibody (αCD4). Selective depletion of NK cells using an anti-

NK1.1 (αNK) antibody in WT compared to (D) MPO inhibitor (MPOi) treated WT mice and (E) HCQ treated WT mice. IgG treated HCQ mice included controls; IgG treated WT and MPOi mice are same dataset as in figure 6. Flow cytometry plots and corresponding quantification confirming depletion of (F) B cells, (G) $CD8^+$ T cells, (H) $CD4^+$ T cells and (I) NK cells from the spleen. Data shown as mean ± SD (n = 4 – 8 mice/group, equal sex); statistical significance of differences calculated using a two-way ANOVA followed by Tukey's multiple comparisons test or unpaired t test (p values shown on graphs).

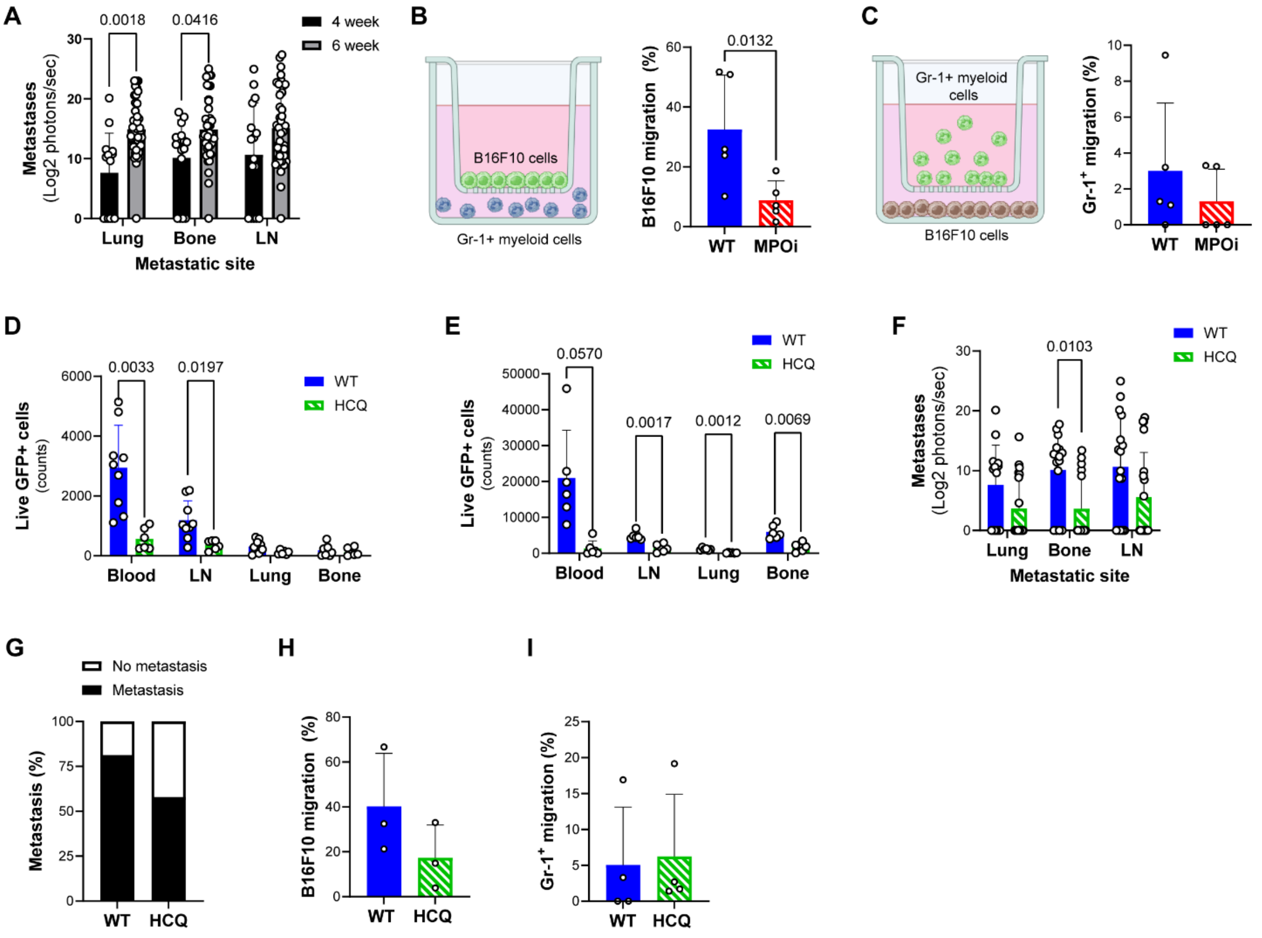


**Figure S10. MPO inhibition decreases B16F10 cell migration and HCQ treatment decreases disseminated tumor cells.** (A) Ex vivo bioluminescence quantification of untreated WT B16F10 spontaneous metastasis mice at 4 weeks compared to 6 weeks. Transwell migration schematic and corresponding quantification of (B) B16F10 migration to Gr-1+ myeloid cells and (C) Gr-1+ myeloid cell migration to B16F10 cells with and without MPO inhibitor (MPOi) treatment (n = 5 independent experiments). Flow cytometry quantification of GFP+ B16F10 cells following (D) a single dose of HCQ at day 19 and (E) 10 days of HCQ treatment at day 28 in the blood, inguinal lymph nodes (LN), lungs and bone of spontaneous B16F10 metastatic WT and HCQ treated mice (n = 6 mice/group; equal sex; WT mice same dataset as in figure 6). (F) *Ex vivo* bioluminescence quantification of metastatic burden and (G) percentage of mice with detectable metastasis at day 28 in WT and HCQ treated mice (n = 19 HCQ; equal sex, WT mice same dataset as in figure 6). Transwell migration of (H) B16F10 cell migration to Gr-1+ myeloid cells and (I) Gr-1+ myeloid cell migration to B16F10 cells with and without HCQ treatment (n = 3 – 4 independent experiments). Data shown as mean ± SD; statistical significance of differences calculated using a two-way ANOVA followed by Tukey's multiple comparisons test, Šídák's multiple comparisons test or paired t test (p values shown on graphs).

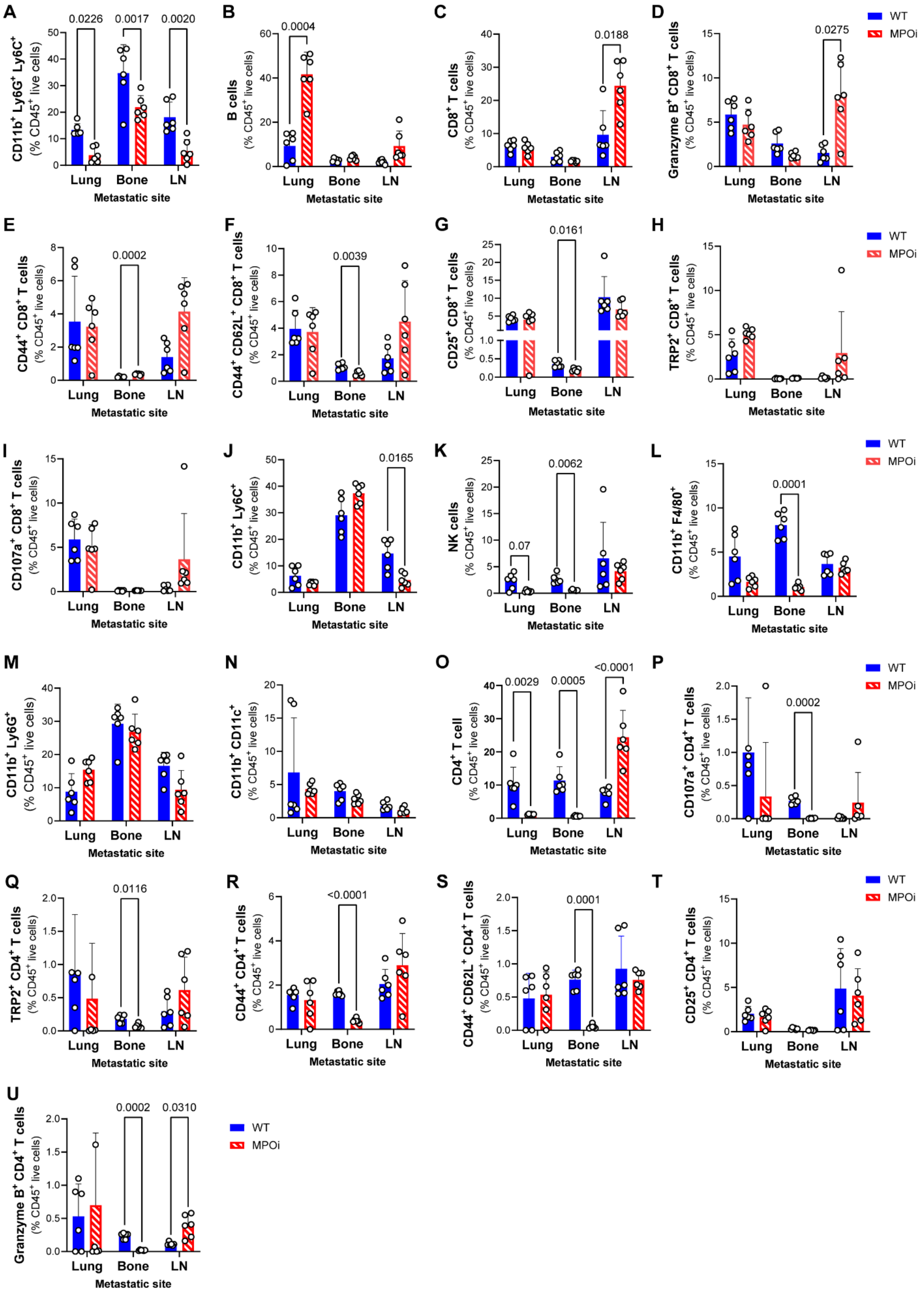

A
CD11b+ Ly6G+ Ly6C+ (% CD45+ live cells)
0.0226
0.0017
0.0020
B
B cells (% CD45+ live cells)
0.0004
C
CD8+ T cells (% CD45+ live cells)
0.0188
D
Granzyme B+ CD8+ T cells (% CD45+ live cells)
0.0275
E
CD44+ CD8+ T cells (% CD45+ live cells)
0.0002
F
CD44+ CD62L+ CD8+ T cells (% CD45+ live cells)
0.0039
G
CD25+ CD8+ T cells (% CD45+ live cells)
0.0161
H
TRP2+ CD8+ T cells (% CD45+ live cells)
I
CD107a+ CD8+ T cells (% CD45+ live cells)
J
CD11b+ Ly6C+ (% CD45+ live cells)
0.0165
K
NK cells (% CD45+ live cells)
0.07
0.0062
L
CD11b+ F4/80+ (% CD45+ live cells)
0.0001
M
CD11b+ Ly6G+ (% CD45+ live cells)
N
CD11b+ CD11c+ (% CD45+ live cells)
O
CD4+ T cell (% CD45+ live cells)
0.0029
0.0005
<0.0001
P
CD107a+ CD4+ T cells (% CD45+ live cells)
0.0002
Q
TRP2+ CD4+ T cells (% CD45+ live cells)
0.0116
R
CD44+ CD4+ T cells (% CD45+ live cells)
<0.0001
S
CD44+ CD62L+ CD4+ T cells (% CD45+ live cells)
0.0001
T
CD25+ CD4+ T cells (% CD45+ live cells)
U
Granzyme B+ CD4+ T cells (% CD45+ live cells)
0.0002
0.0310
Lung
Bone
LN
Metastatic site
WT
MPOi

**Figure S11. MPO inhibition remodels metastatic microenvironment at an earlier timepoint.** Flow cytometry quantification of frequencies (%) of (A) $Ly6G^{+}Ly6C^{+}$ myeloid cells, (B) B cells, (C) $CD8^{+}$ T cells, (D) $GranzymeB^{+}$ $CD8^{+}$ T cells, (E) $CD44^{+}$ $CD8^{+}$ T cells, (F) $CD44^{+}CD62L^{+}$ $CD8^{+}$ T cells, (G) $CD25^{+}$ $CD8^{+}$ T cells, (H) $TRP2^{+}$ $CD8^{+}$ T cells, (I) $CD107a^{+}$ $CD8^{+}$ T cells, (J) $Ly6C^{+}$ monocytes, (K) NK cells, (L) $F4/80^{+}$ macrophages, (M) $Ly6G^{+}$ neutrophils, (N) $CD11b^{+}CD11c^{+}$ dendritic cells, (O) $CD4^{+}$ T cells, (P) $CD107a^{+}$ $CD4^{+}$ T cells, (Q) $TRP2^{+}$ $CD4^{+}$ T cells, (R) $CD44^{+}$ $CD4^{+}$ T cells, (S) $CD44^{+}CD62L^{+}$ $CD4^{+}$ T cells, (T) $CD25^{+}$ $CD4^{+}$ T cells, (U) $GranzymeB^{+}$ $CD4^{+}$ T cells in the lungs, bone, and inguinal lymph nodes (LN) from spontaneous B16F10 metastatic WT and MPO inhibitor treated (MPOi) WT mice at 28 days. Data shown as mean ± SD (n = 6 mice/group, equal sex); statistical significance of differences in immune cell frequencies calculated using a two-way ANOVA followed by Šídák's multiple comparisons test (p values shown on graphs).

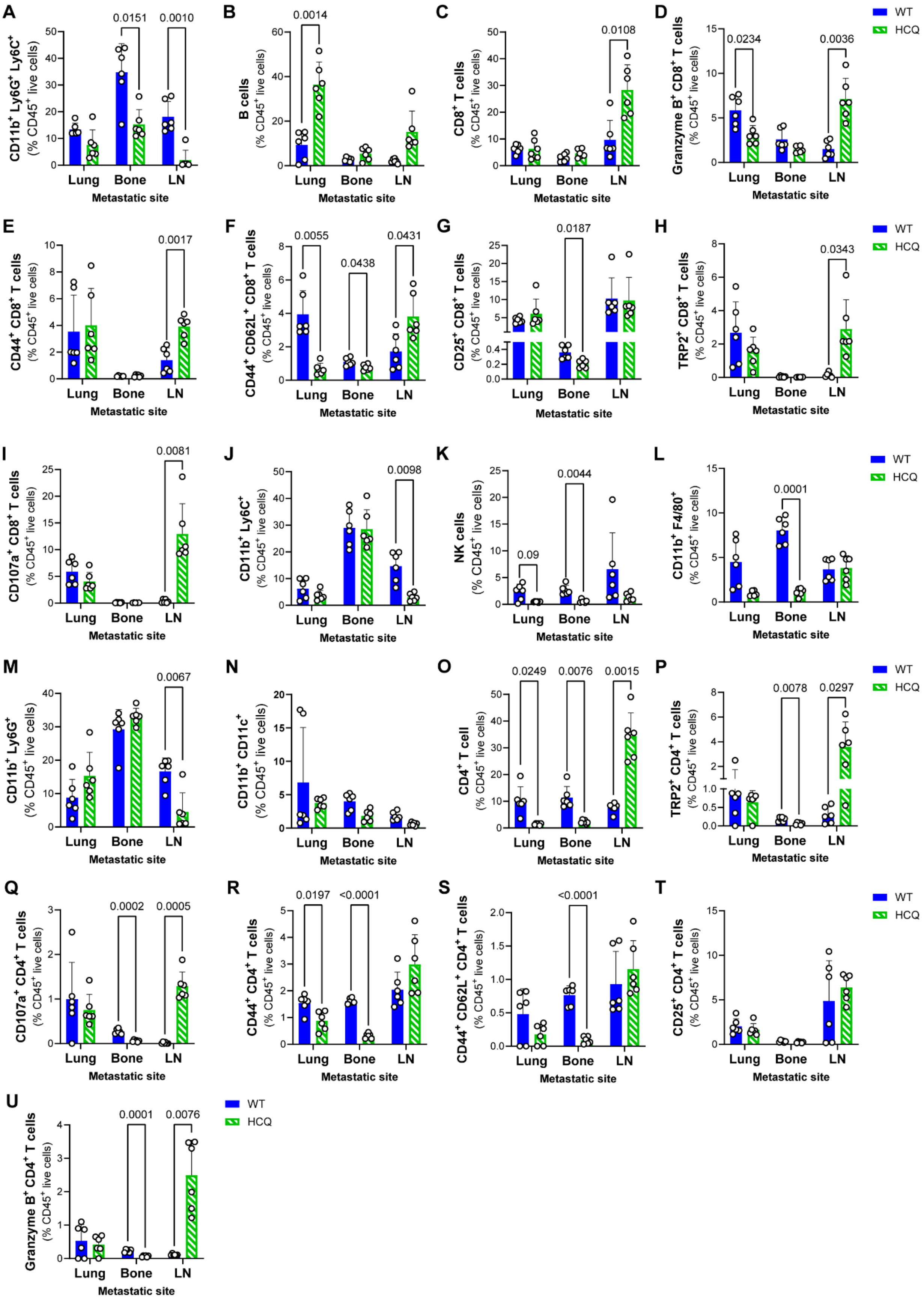

A
CD11b+ Ly6G+ Ly6C+ (% CD45+ live cells)
0.0151
0.0010
B
B cells (% CD45+ live cells)
0.0014
C
CD8+ T cells (% CD45+ live cells)
0.0108
D
Granzyme B+ CD8+ T cells (% CD45+ live cells)
0.0234
0.0036
E
CD44+ CD8+ T cells (% CD45+ live cells)
0.0017
F
CD44+ CD62L+ CD8+ T cells (% CD45+ live cells)
0.0055
0.0438
0.0431
G
CD25+ CD8+ T cells (% CD45+ live cells)
0.0187
H
TRP2+ CD8+ T cells (% CD45+ live cells)
0.0343
I
CD107a+ CD8+ T cells (% CD45+ live cells)
0.0081
J
CD11b+ Ly6C+ (% CD45+ live cells)
0.0098
K
NK cells (% CD45+ live cells)
0.09
0.0044
L
CD11b+ F4/80+ (% CD45+ live cells)
0.0001
M
CD11b+ Ly6G+ (% CD45+ live cells)
0.0067
N
CD11b+ CD11c+ (% CD45+ live cells)
O
CD4+ T cell (% CD45+ live cells)
0.0249
0.0076
0.0015
P
TRP2+ CD4+ T cells (% CD45+ live cells)
0.0078
0.0297
Q
CD107a+ CD4+ T cells (% CD45+ live cells)
0.0002
0.0005
R
CD44+ CD4+ T cells (% CD45+ live cells)
0.0197
<0.0001
S
CD44+ CD62L+ CD4+ T cells (% CD45+ live cells)
<0.0001
T
CD25+ CD4+ T cells (% CD45+ live cells)
U
Granzyme B+ CD4+ T cells (% CD45+ live cells)
0.0001
0.0076
Lung
Bone
LN
Metastatic site
WT
HCQ

**Figure S12. MPO inhibition remodels metastatic microenvironment at an earlier timepoint.** Flow cytometry quantification of frequencies (%) of (A) $Ly6G^+Ly6C^+$ myeloid cells, (B) B cells, (C) $CD8^+$ T cells, (D) $GranzymeB^+$ $CD8^+$ T cells, (E) $CD44^+$ $CD8^+$ T cells, (F) $CD44^+CD62L^+$ $CD8^+$ T cells, (G) $CD25^+$ $CD8^+$ T cells, (H) $TRP2^+$ $CD8^+$ T cells, (I) $CD107a^+$ $CD8^+$ T cells, (J) $Ly6C^+$ monocytes, (K) NK cells, (L) $F4/80^+$ macrophages, (M) $Ly6G^+$ neutrophils, (N) $CD11b^+CD11c^+$ dendritic cells, (O) $CD4^+$ T cells, (P) $CD107a^+$ $CD4^+$ T cells, (Q) $TRP2^+$ $CD4^+$ T cells, (R) $CD44^+$ $CD4^+$ T cells, (S) $CD44^+CD62L^+$ $CD4^+$ T cells, (T) $CD25^+$ $CD4^+$ T cells, (U) $GranzymeB^+$ $CD4^+$ T cells in the lungs, bone, and inguinal lymph nodes (LN) from spontaneous B16F10 metastatic WT and HCQ treated WT mice at 28 days. Data shown as mean ± SD (n = 6 mice/group, equal sex; WT dataset same as in figure S11); statistical significance of differences in immune cell frequencies calculated using a two-way ANOVA followed by Šídák's multiple comparisons test (p values shown on graphs).

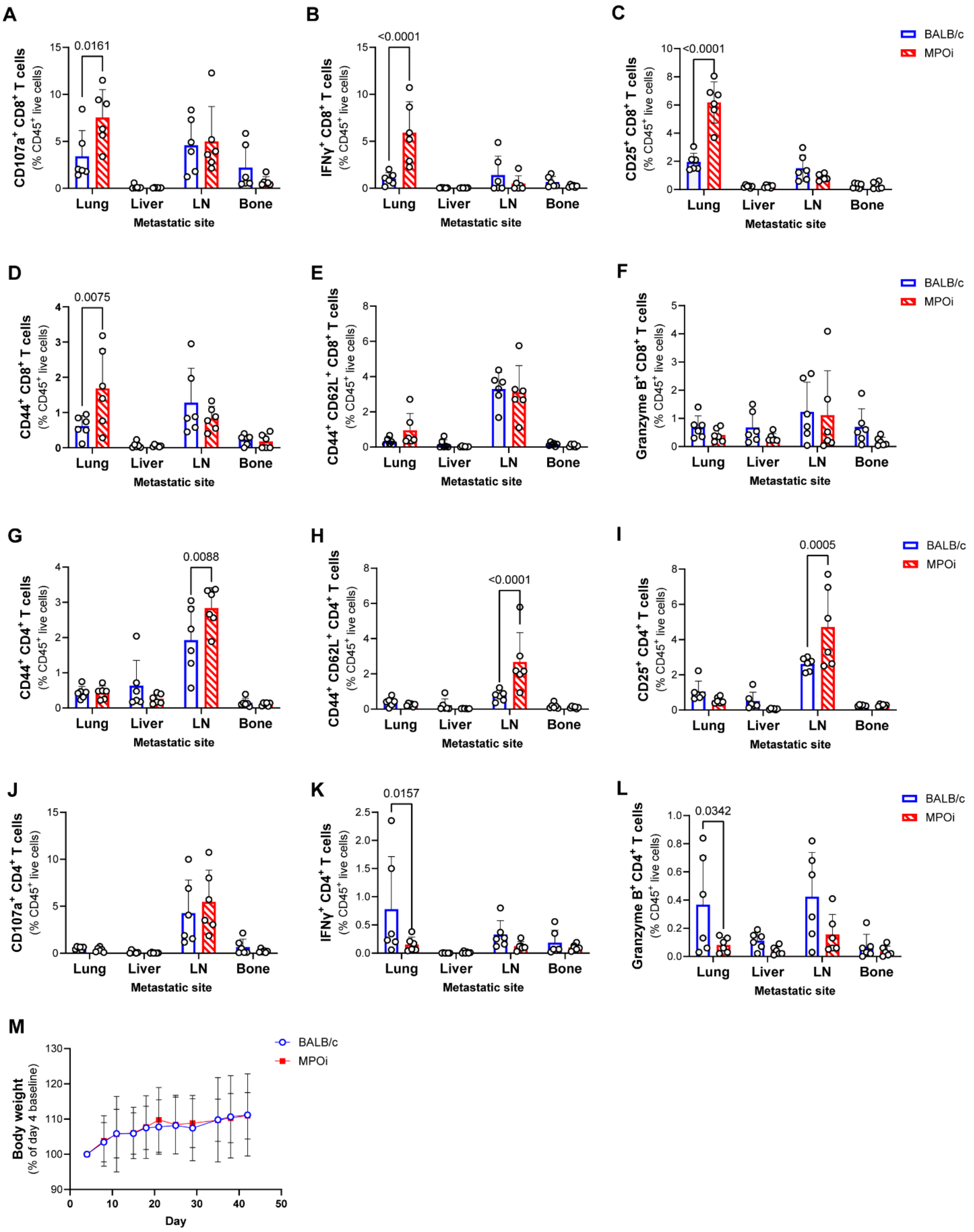


**Figure S13. MPO inhibition enhances the frequency of activated and effector CD8+ T cells in metastatic breast cancer related to Figure 7.** Flow cytometry quantification of frequencies (%) of (A) CD107a+ CD8+ T cells, (B) IFNγ+ CD8+ T cells, (C) CD25+ CD8+ T cells, (D) CD44+

$CD8^+$ T cells, (E) $CD44^+CD62L^+$ $CD8^+$ T cells, (F) $GranzymeB^+$ $CD8^+$ T cells, (G) $CD44^+$ $CD4^+$ T cells, (H) $CD44^+CD62L^+$ $CD4^+$ T cells, (I) $CD25^+$ $CD4^+$ T cells, (J) $CD107a^+$ $CD4^+$ T cells, (K) $IFN\gamma^+$ $CD4^+$ T cells, (L) $GranzymeB^+$ $CD4^+$ T cells in the lungs, liver, inguinal lymph nodes (LN) and bone from spontaneous 4T-1 metastatic BALB/c and MPO inhibitor treated (MPOi) BALB/c mice at experimental endpoint (n = 6 female mice/group). (M) Body weight measurements of BALB/c and MPOi mice (n = 8 female mice/group). Data shown as mean ± SD; statistical significance of differences in immune cell frequencies calculated using a two-way ANOVA followed by Šídák's multiple comparisons test (p values shown on graphs).

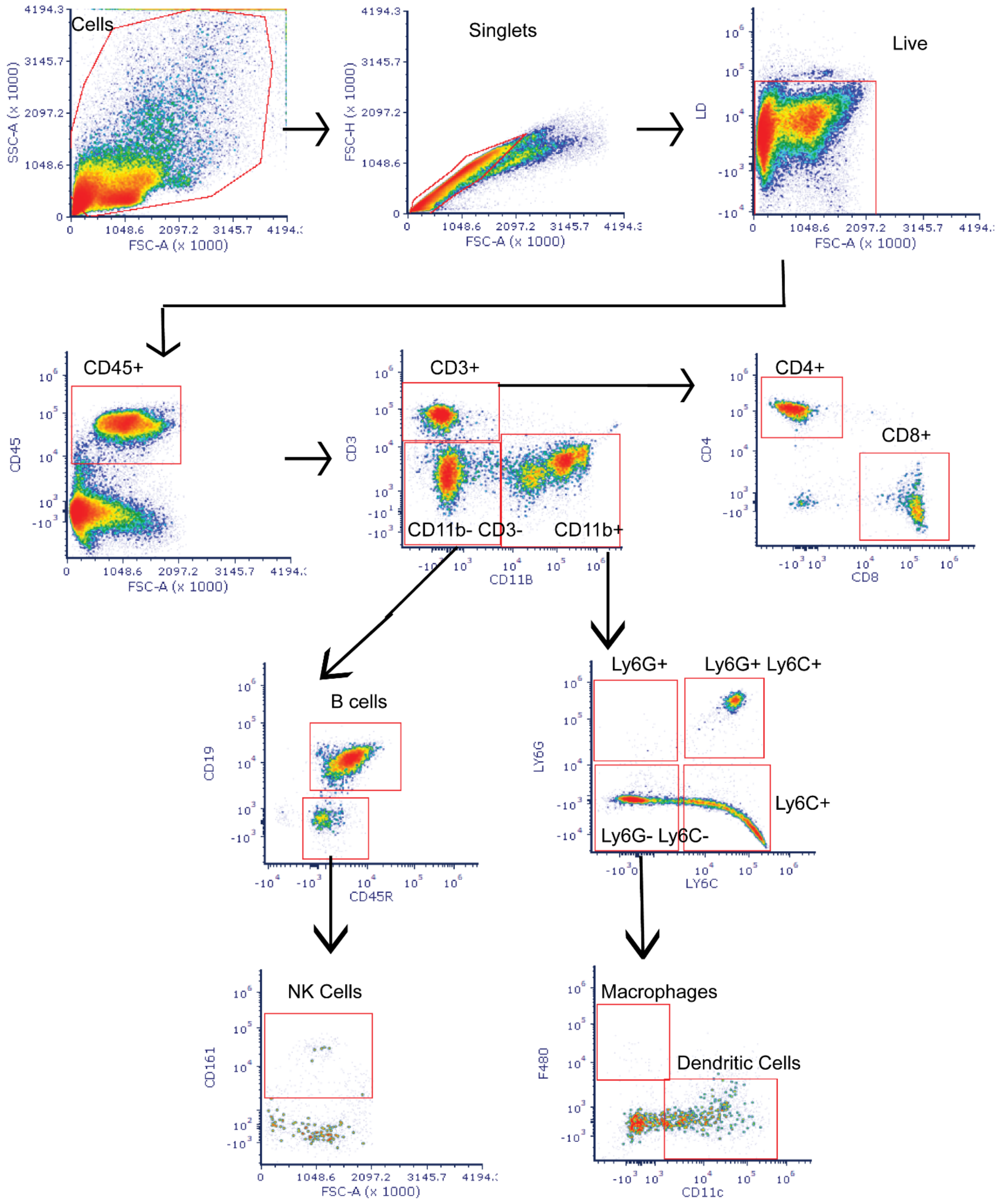


**Figure S14. Flow cytometry immune cell panel gating strategy.**

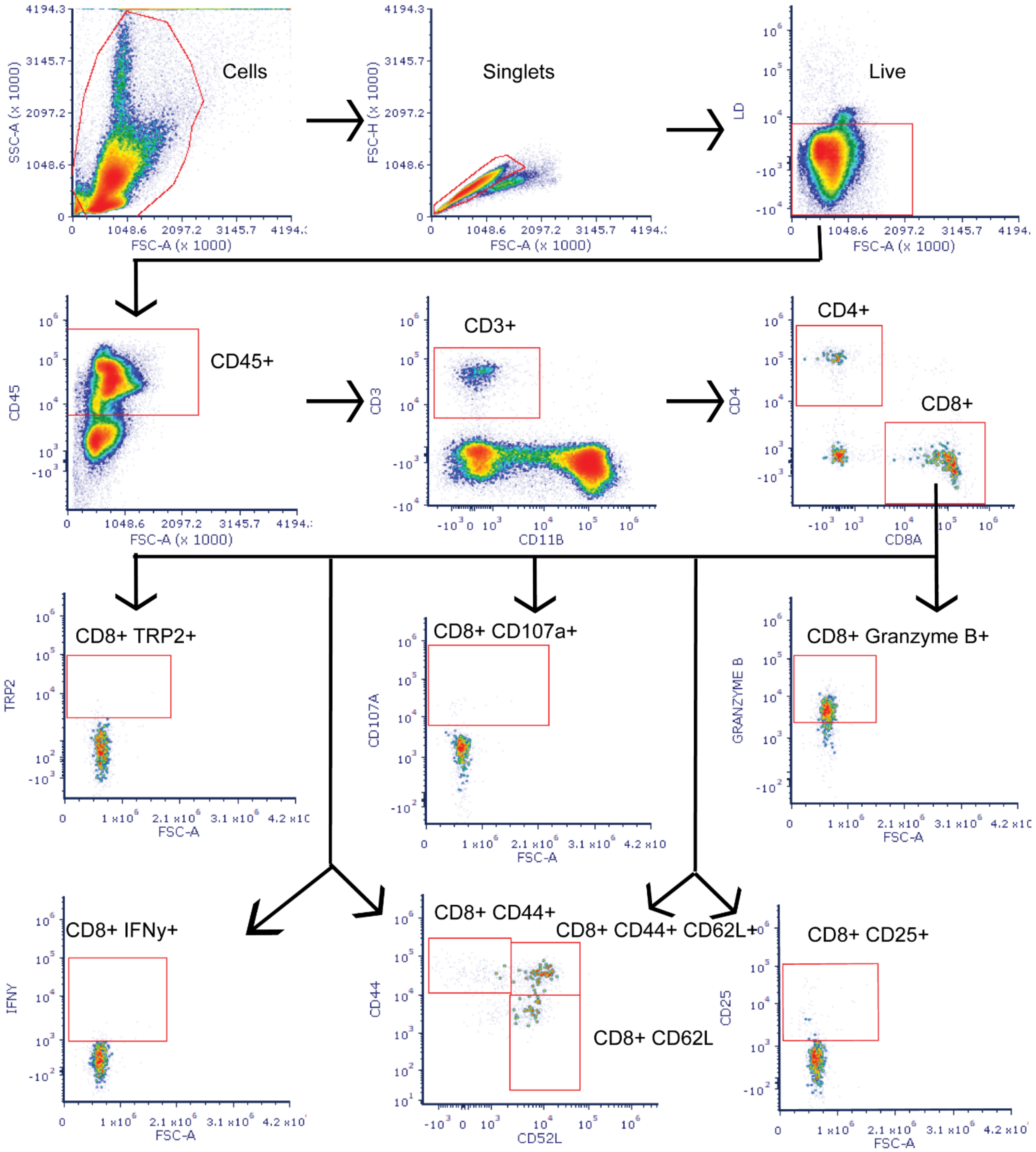


**Figure S15. Flow cytometry CD8$^+$ T cell panel gating strategy.**

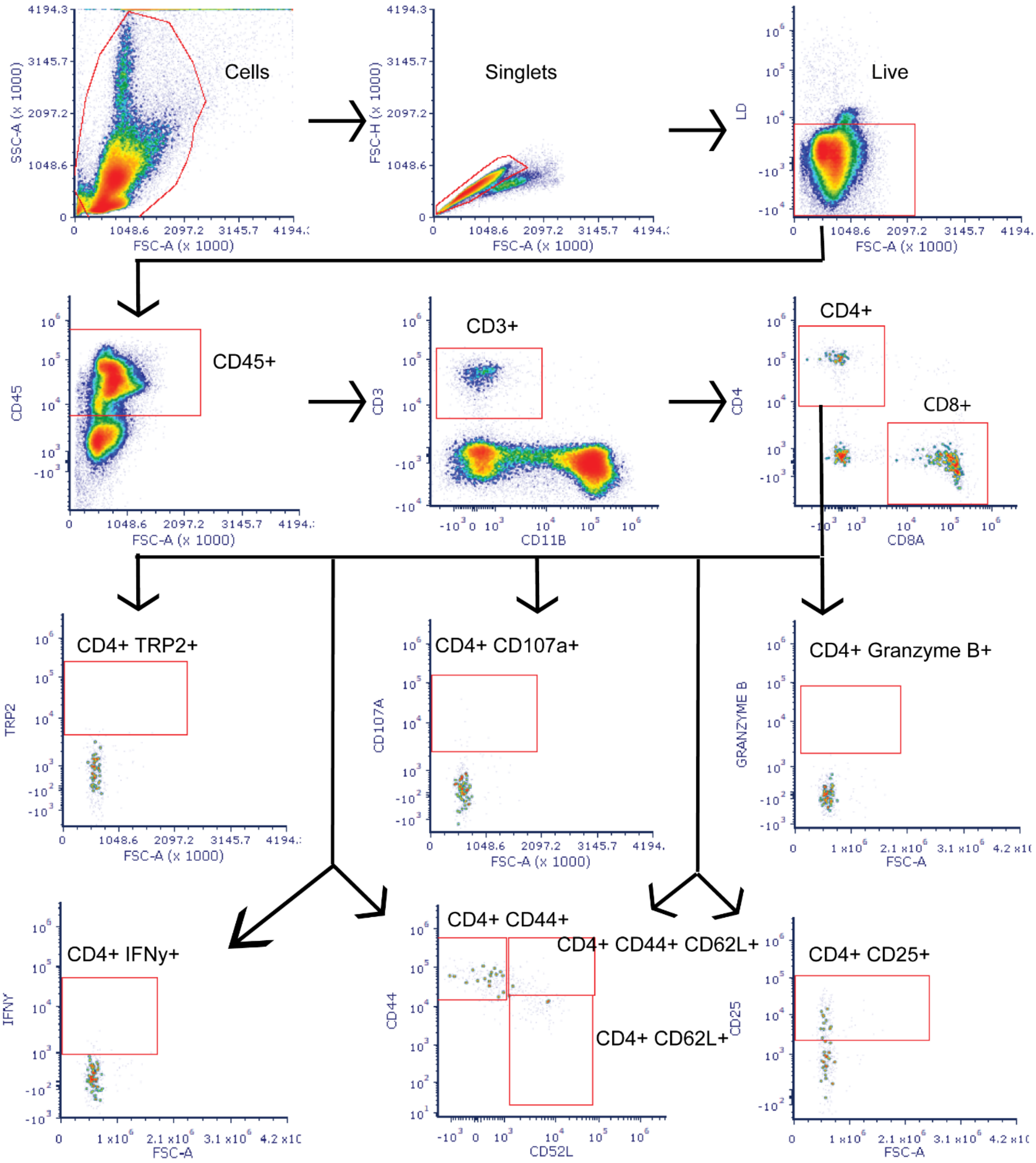


**Figure S16. Flow cytometry CD4$^+$ T cell panel gating strategy.**

**Table 1. Flow cytometry immune cell antibody list**

| Marker | Fluorophore | Company |
|---|---|---|
| CD3 | Alexa Fluor 488 | BioLegend (CA, USA) |
| CD8a | APC-Fire 750 | BioLegend (CA, USA) |
| CD19 | APC-Fire 810 | BioLegend (CA, USA) |
| F4/80 | Brilliant Violet 605 | BioLegend (CA, USA) |
| CD11c | Brilliant Violet 421 | BioLegend (CA, USA) |
| CD161 | Brilliant Violet 711 | BioLegend (CA, USA) |
| CD4 | Brilliant Violet 785 | BioLegend (CA, USA) |
| Ly-6C | PerCP | BioLegend (CA, USA) |
| CD45 | Pacific Blue | BioLegend (CA, USA) |
| Ly-6G | PE-Fire 640 | BioLegend (CA, USA) |
| CD45R | Spark Blue 550 | BioLegend (CA, USA) |
| CD11b | Spark NIR 685 | BioLegend (CA, USA) |
| Fixable Live/Dead | Zombie Aqua | BioLegend (CA, USA) |
| CD25 | APC | BioLegend (CA, USA) |

**Table 2. Flow cytometry T cell antibody list**

| Antibody | Fluorophore | Company |
|---|---|---|
| CD3 | Alexa Fluor 488 | BioLegend (CA, USA) |
| CD11b | Spark NIR 685 | BioLegend (CA, USA) |
| CD8a | APC-Fire 750 | BioLegend (CA, USA) |
| CD4 | Brilliant Violet 785 | BioLegend (CA, USA) |
| CD25 | PE-Cy7 | BioLegend (CA, USA) |
| PD-1 | PE-Dazzle 594 | BioLegend (CA, USA) |
| CTLA-4 | Brilliant Violet 421 | BioLegend (CA, USA) |
| Granzyme B | APC | BioLegend (CA, USA) |
| CD44 | Brilliant Violet 570 | BioLegend (CA, USA) |
| CD62L | Brilliant Violet 650 | BioLegend (CA, USA) |
| TRP2 | PE | Santa Cruz Biotechnology (TX, USA |
| CD45 | Pacific Blue | BioLegend (CA, USA) |
| IFN-gamma | PerCP-Cy5.5 | BioLegend (CA, USA) |
| Fixable Live/Dead | Zombie Aqua | BioLegend (CA, USA) |
| CD107a | Alexa Fluor 700 | BioLegend (CA, USA) |
